# Cell dichotomous role of STING in pulmonary hypertension

**DOI:** 10.1101/2022.11.29.518422

**Authors:** Ann T. Pham, Aline C. Oliveira, Chunhua Fu, Matthew D. Alves, Zadia Dupee, Laylo Mukhsinova, Elnaz Ebrahimi, Harsh Patel, Reeha Patel, Amy Nguyen, Lei Jin, Andrew J. Bryant

## Abstract

**Rationale:** Patients with constitutive activation of DNA sensing pathway through stimulator of interferon genes (STING), such as those with STING-Associated Vasculopathy with onset in Infancy (SAVI), frequently have complications related to pulmonary hypertension (PH). However, the role of STING-signaling in adult PH patients is heretofore undescribed.

**Objective:** To investigate the role of STING in PH development.

**Methods and Results:** PH was induced in global STING deficient or cell-specific STING deficient mice using either bleomycin or chronic hypoxia exposure. PH development was evaluated with right ventricular systolic pressure, Fulton index, histological and flow cytometric measurements. STING expression in patient lungs were examined using both immunohistochemistry and flow cytometry. Herein, we describe how STING overactivation in a SAVI mouse model results in a baseline elevation in pulmonary pressures, while global STING deficiency protects mice from PH development. Furthermore, STING-associated PH appears to be independent of type I Interferon (IFN) signaling. We further demonstrate a cellular dichotomous role of STING in PH development with STING expression by smooth muscle cells contributing to PH, and its activation on myeloid cells being pivotal in severe disease prevention. Finally, we demonstrate a STING-PD-L1 axis as necessary for disease progression, suggesting future potential therapeutic applications.

**Conclusions:** Overall, these data provide concrete evidence of STING involvement in PH, establishing biologic plausibility for STING-related therapies in PH treatment.

**Graphical abstract:** 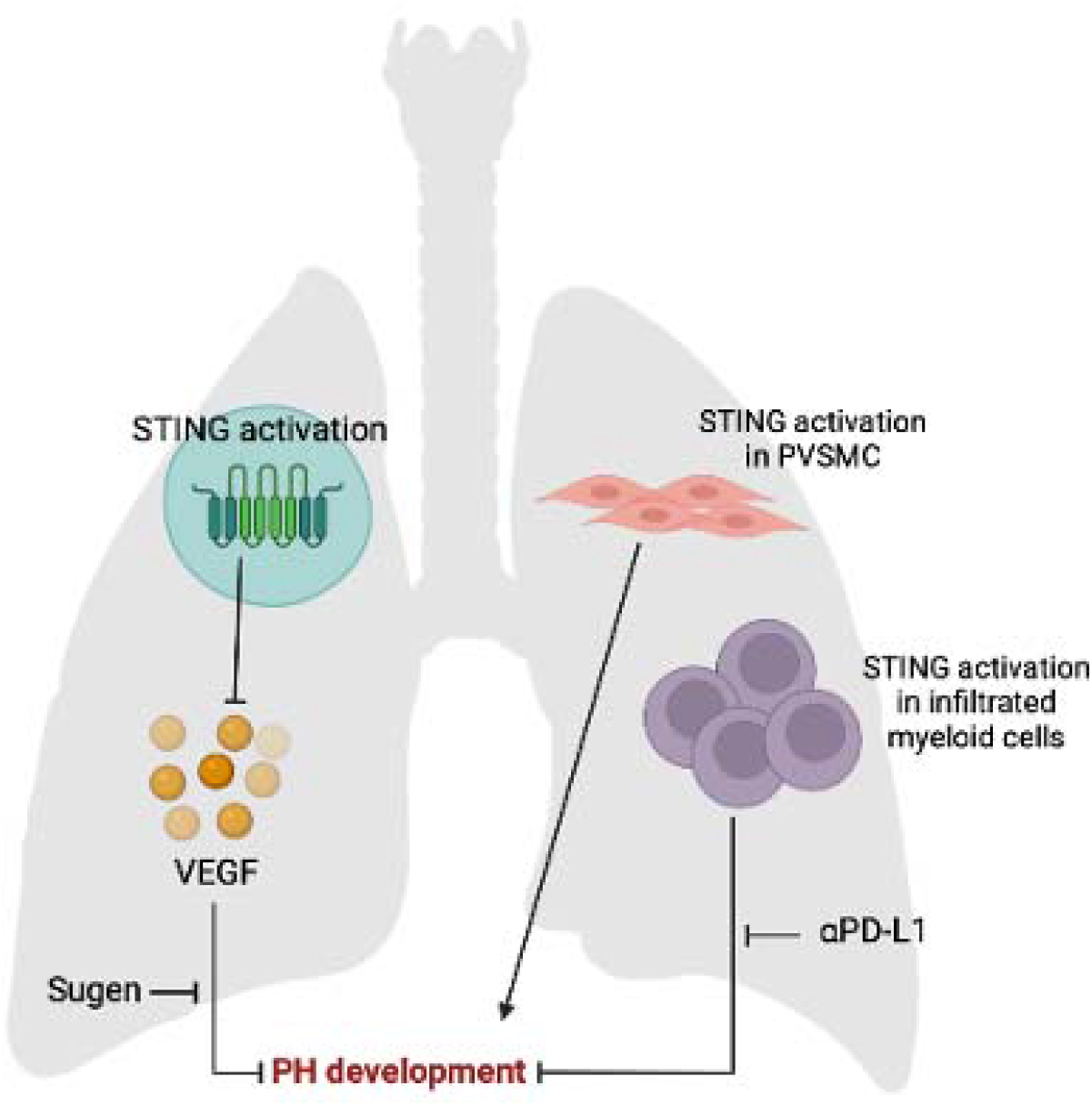

## Introduction

Pulmonary hypertension (PH) is a devastating disease, affecting approximately 15-50 million people worldwide^1^. World Health Organization (WHO) Group 3 PH – PH secondary to chronic lung disease and/or hypoxia – is the second leading cause of PH, with rising incidence globally and the highest mortality rate among all types of pulmonary vascular disease^2–4^. Group 3 PH patients, likewise, utilize the health care system up to four times higher than patients without complicating PH^5^. There is a significant therapeutic gap for afflicted patients, with currently only one PH-specific drug indicated for use in patients^6^. More research is thus needed into understanding disease pathogenesis.

Previous studies have demonstrated a critical role for immune cells in disease progression^7,8^. Enhanced discernment of the various immune cell subtypes in PH represents a novel area of research into disease modifying therapy. In patients with genetic predisposition to constitutive interferon production, a family of illnesses referred to as “interferonopathies”, there is a proclivity to PH development^9–13^. Such disease states provide a novel link between systemic inflammation and pulmonary vascular remodeling. The canonical interferonopathy that predisposes to development of pulmonary vasculopathy, termed STING-Associated Vasculopathy onset in Infancy (SAVI), is observed in children sharing activating mutations in the upstream regulator of type I IFN signaling, Stimulator of Interferon Genes (STING)^12,13^. STING is a cellular gatekeeper against a variety of cytosolic DNA-containing pathogens ^14^, serving to detect and respond to – evolutionarily – viral DNA. Upon activation of STING, a series of downstream transcription factors, including Interferon regulator factor 3 (IRF3), are subsequently activated. Phosphorylated IRF3 enters the nucleus and promotes transcription of type I IFN genes and interferon-response genes^14,15^, resulting in immune cell activation. Notably, a recent study of STING pathway analysis in COVID-19 revealed a pathogenic role for STING, contributing to lung inflammation and disease progression^16^. Likewise, STING promotes inflammation in a variety of disparate disease models^17–19^, playing an important role in immune homeostasis^20^, contributing in particular to parenchymal lung disease and systemic vasculitis^21^. To date, there are no data on STING involvement in pulmonary vascular disease, although the role of type I IFN in PH have been addressed by different research groups, albeit with contradicting results^22,23^. As PH has been associated with DNA damage due to drug and toxin-mediated circulating free DNA^24^, and an increase level in circulating DNA due to predisposing disease-associated mutations^25,26^, it makes intuitive sense that downstream detection of free DNA, through STING, may contribute to disease.

Herein, we report on the relevance of STING in patients with interstitial lung disease. Next, we evaluate pulmonary vascular response in a validated SAVI mouse model. We go on to describe that global STING deficient (STING^-/-^) mice are protected against PH in two complimentary models of disease, with phenotype developing independent of interferon up-regulation. We then describe how STING regulates PH development and progression in a cell-specific manner. Specifically, we demonstrate that smooth muscle expression of STING contributes to disease pathogenesis, and STING expression on myeloid cells is necessary to prevent development of severe PH. Finally, we show an important yet complex role for STING-VEGF signaling in protection against PH. Together, our data not only support a role for STING in PH development but lays the groundwork for STING-based therapies that can be utilized in PH patients^27^.

## Results

### Patients with PH-predisposing lung disease display increased pulmonary STING expression

To investigate STING application in World Health Organization (WHO) Group 3 PH, that is secondary to parenchymal lung disease or chronic hypoxia, we examined STING expression grossly in patients with interstitial lung disease (ILD), as well as patients with PH and associated pulmonary disease, namely idiopathic pulmonary fibrosis (IPF). To this end, lung single-cell suspensions from healthy individuals and patients diagnosed with ILD (n=3/group) were subjected to flow cytometry for analysis of STING expression in various cell groups. Notably, there was an increase in STING expression in CD33^+^CD11b^+^HLA^-^DR^-^CD14^-^CD15^+^ cells, previously described as human polymorphonuclear myeloid-derived suppressor cells (PMN-MDSC)^28^ (**Fig. 1, A – C**). The equivalent cell population in mice is the pathogenic CD11b^+^Ly6C^lo^Ly6G^+^, to which we previously demonstrated a pathogenic role in PH development^29^. As PH is a known common comorbidity for ILD, we were strongly encouraged to move forward with our hypothesis of STING involvement in associated-PH. We then examined formalin-fixed lung sections of IPF patients with and without PH, in a previously defined cohort^30^, and found a purely qualitative abundance of STING expression in the lung sections of patients with IPF and IPF+PH compared to healthy individuals (**Fig. 1D**). Although we were unable to directly compare STING expression through quantification between IPF and IPF+PH groups, we noted high levels of STING expression in IPF+PH patients, as well (**Fig. 1D**). Similarly, chronic obstructive pulmonary disease (COPD) with PH patients also demonstrated elevated level of pulmonary STING expression, seemingly more so than COPD patients and healthy individuals (**Fig. S1**). These results, together with observations in SAVI patients and mouse model, further highlighted the potential relevance of STING to PH development, emboldening further investigation.

**Figure 1:**
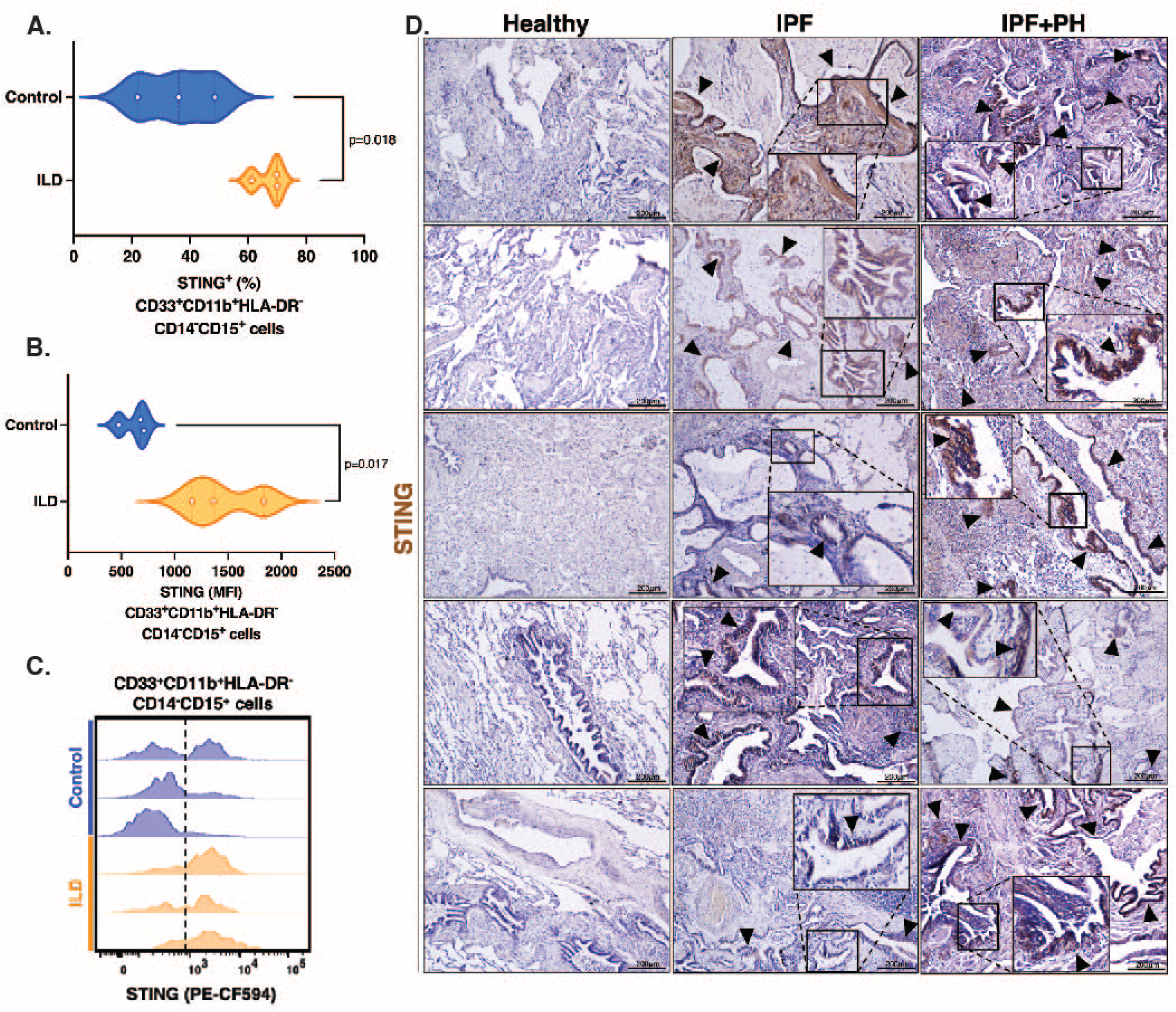
Patients with PH-predisposing lung disease display increased pulmonary STING expression. (**A**) % Expression, (**B**) MFI quantification and (**C**) Representative flow plots of STING expression in pulmonary infiltrated CD33^+^CD11b^+^HLA^-^DR^-^CD14^-^CD15^+^ cells of healthy and interstitial lung disease (ILD) patients. Each dot presents an individual donor. Level of significance was calculated with unpaired two-tail student’s T test. P<0.05 was considered significant. (**D**) Representative images of immunohistochemical (IHC) staining of STING (brown, arrowheads) in formalin-fixed lung sections of healthy individuals and patients with idiopathic pulmonary fibrosis (IPF) with or without PH at 10x magnification. Scale bar = 200mm. Each image is from an individual donor (n=5/group).

### Global STING deficient mice are protected against PH development secondary to bleomycin and chronic hypoxia

Since STING appears to be expressed in patients with chronic parenchymal lung disease, we hypothesized that deletion of STING would protect against PH development. To that end, we induced PH in 8–10-week-old global STING deficient (STING^-/-^) mice using either bleomycin or chronic hypoxia as previously described^29,31^. Deletion of STING was confirmed with both genotyping and western blotting of whole lung protein (**Fig. S2A**). Upon assessment of pulmonary hemodynamics, bleomycin- and chronic hypoxia-induced PH control mice were found to have elevated right ventricular systolic pressure (RVSP), while STING^-/-^ mice displayed significantly lower RVSP (**Fig. 2, A and D**). Surprisingly, right ventricular (RV) remodeling was not observed in STING^-/-^ mice exposed to chronic hypoxia, assessed by Fulton Index (ratio of right ventricular mass over left ventricular and septum mass), in contrast to chronic hypoxic control mice, which displayed an increase in RV mass by percentage (**Fig. 2B**). a-smooth muscle actin (aSMA) immunohistochemical (IHC) staining revealed a trend in attenuation of complete muscularized vessels as well as total muscularized pulmonary vessels, especially in the large and small vessels, in chronic hypoxia STING^-/-^ mice (**Figure 2, G – F**). As bleomycin induces PH presumably through a fibrosis-related mechanism, we performed Masson Trichrome (MTC) staining on control and bleomycin treated groups to assess degree of semi-quantitative fibrotic changes. To that end, inflammation/fibrosis assessment with MTC staining showed an anticipated decrease in inflammation in the lung of STING^-/-^ mice treated with bleomycin compared to control (**Fig. 2, G and H**). Collectively, these data indicate that deletion of STING provides protection against elevation in pulmonary pressure.

**Figure 2.**
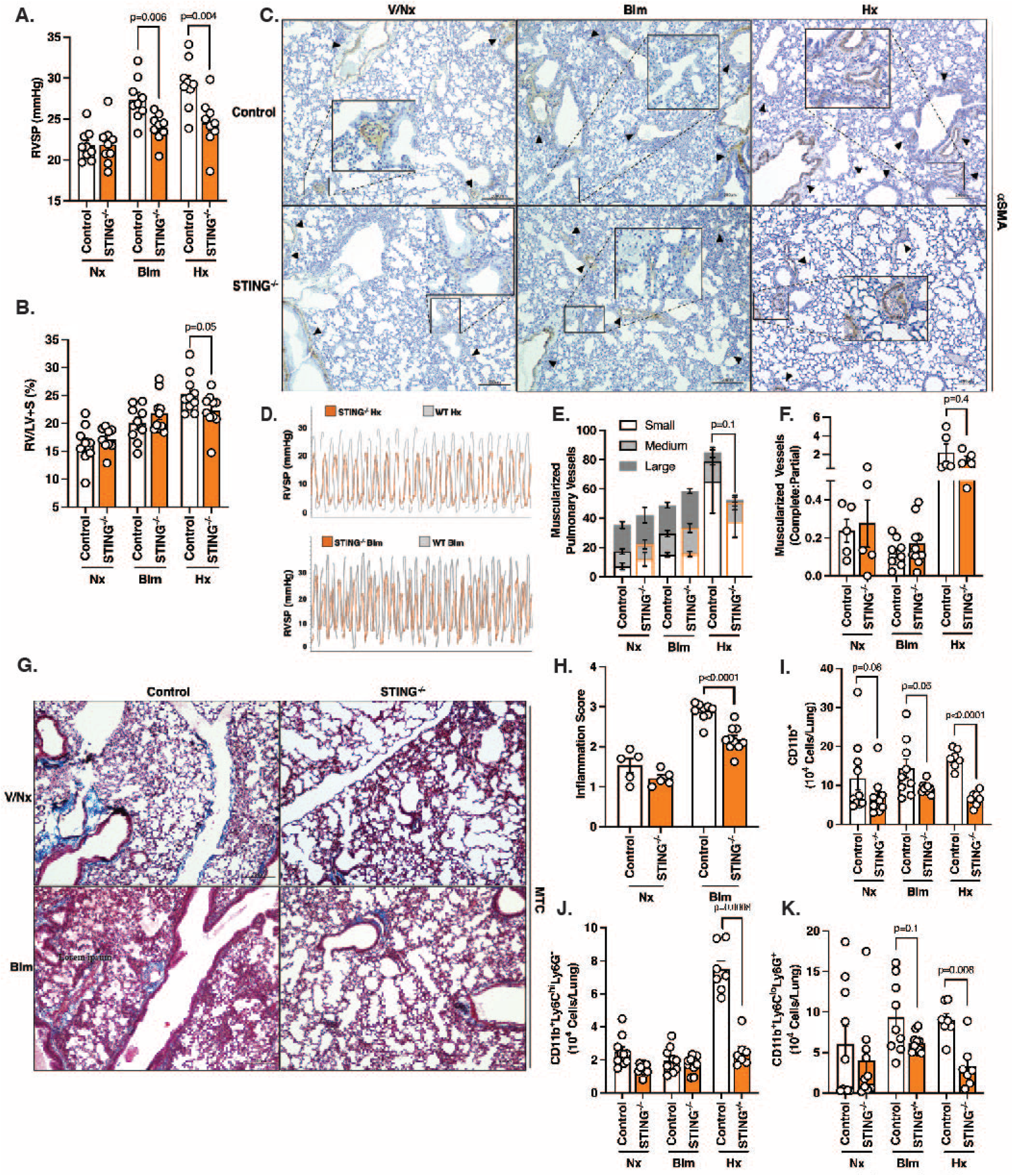
Global STING deficient mice are protected against PH development secondary to bleomycin and chronic hypoxia. (**A**) Invasive right ventricular systolic pressure (RVSP) measurements and (**D**) curve representations of wild type (WT) control and global STING deficient (STING^-/-^) mice subjected to normoxia (Nx), bleomycin (Blm) or chronic hypoxia (Hx). (**B**) Fulton Index of WT and STING^-/-^ mice from designated groups. (**C**) Representative images of a-smooth muscle actin (aSMA, brown, arrowheads) IHC staining on formalin-fixed lung sections of dedicated mouse groups. Scale bar = 200mm at 10x magnification. (**E and F**) Quantification of muscularized pulmonary vessels (small, medium, large, complete and partial) from IHC staining of aSMA of mice from all groups. (**G**) Representative images of Masson Trichrome (MTC)-stained formalin-fixed lung sections of control and bleomycin treated mice. Scale bar = 200mm at 10x magnification. (**H**) Inflammation and fibrosis scoring from MTC-stained formalin-fixed lung sections of WT and STING^-/-^ mice undergoing bleomycin treatment. (**I** – **K**) Flow cytometric quantification of pulmonary infiltrated (**I**) CD11b^+^, (**J**) CD11b^+^Ly6C^hi^L6G^-^, and (**K**) CD11b^+^Ly6^lo^Ly6G^+^ cells of WT and STING^-/-^ mice subjected to normoxia, bleomycin, or chronic hypoxia. Cell identity gates are shown in **Fig. 3K**. Each dot is an individual mouse (n=5-8/group). Column represents mean ± SEM. Significance level was calculated with un-paired two-tailed student T’s test. P<0.05 was considered significant.

We have previously demonstrated the important role of myeloid cells, especially myeloid-derived suppressor cells (MDSCs), in PH and related pulmonary disease^29–32^. While the context and classification of MDSCs are still evolving, they are commonly described as immature myeloid cells that are morphologically similar to either monocytes (Mo-MDSC, CD11b^+^Ly6C^hi^Ly6G^-^) or neutrophils (PMN-MDSC, CD11b^+^Ly6C^lo^ Ly6G^+^), capable of suppressing other immune cells, especially T cells^28^. As STING is an important regulator of both innate and adaptive immunity, we expected to see changes in pulmonary infiltrated myeloid cells in STING^-/-^ mice, correlating with RVSP changes. Indeed, under both bleomycin treatment and chronic hypoxia exposure, there was a decreased in number of pulmonary infiltrated CD11b^+^ cells in STING^-/-^ mice compared to controls (**Fig. 2I**). Interestingly, STING^-/-^ mice also displayed decreased CD11b^+^ cells under baseline conditions (**Fig. 2I**), which we attributed to decreased inflammatory signaling consistent with loss of STING. We also observed a similar trend of decreased pulmonary infiltrated subpopulations of myeloid cells in PH-induced STING^-/-^ mice, including aforementioned CD11b^+^Ly6C^hi^Ly6G^-^ and CD11b^+^Ly6C^lo^Ly6G^+^ cells (**Fig. 2, J and K**). The difference was more profound in our chronic hypoxia-induced PH model, while no significant difference was found between CD11b^+^Ly6C^hi^Ly6G^-^ of bleomycin treated STING^-/-^ mice and control (**Fig. 2J**). Similarly, although there was not a significant decrease in CD11b^+^Ly6C^lo^Ly6G^+^ population of bleomycin treated STING^-/-^ mice, we detected a trend toward a reduction in their number (**Fig. 2K**). It is worth noting that STING^-/-^ mice exhibited a small population of CD11b^+^Ly6C^lo^Ly6G^-^ cells, most likely composed of alveolar macrophages. Although initial analysis of this cell population yielded no change, we cannot rule out their involvement in STING-mediated PH development based on these data.

To further establish STING relevancy in pulmonary disease, we examined PH development in SAVI mice with N153S mutation, as PH evaluation has never been ascertained in SAVI mouse models. Here, we found that 10-week-old SAVI male mice (N153S mutation) at baseline displayed elevated RVSP compared to wild type (WT) control (**Fig. S3A**). In the contrary, SAVI female mice did not demonstrate a difference in RVSP compared to WT (**Fig. S3A**). Interestingly, although there was not a significant change in the Fulton Index between WT and SAVI male mice, it was not the case for SAVI female mice, indicating RV remodeling, as a consequence of PH (**Fig. S3D**). In addition, MTC staining revealed a significantly higher level of tissue fibrosis in lung sections of both male and female SAVI mice, suggesting a proinflammatory microenvironment (**Fig. S3, C and D**). Indeed, as expected, when looking at the pulmonary immune cell profile of SAVI mice, we saw a rise in number of infiltrated CD11b^+^ cells in SAVI male mice (**Fig. S3E**), including the pathogenic CD11b^+^Ly6C^lo^Ly6G^+^ sub-population (**Fig. S3G**). Similar to our previous findings utilizing different PH mouse models, CD11b^+^Ly6C^hi^Ly6G^-^ cell numbers remained unchanged (**Fig. S3F**). In addition, splenic MDSCs isolated from SAVI male mice exhibited significantly higher immunosuppressive capability toward cytotoxic T cells, namely CD4^+^ and CD8^+^ T cells, correlating with disease development (**Fig. S3, H and I**). Consistent with SAVI mice having constitutive activation of STING, we found an increase in expression of pSTAT3, a downstream regulator of STING-associated type I IFN signaling, in CD11b^+^ and CD11b^+^Ly6C^lo^Ly6G^+^ cells of SAVI male mice (**Fig. S3, J and K**). Together, these findings demonstrate SAVI mice with N153S mutation are susceptible to PH, phenocopying aspects of patients with disease, demonstrating additional robust relevance of STING and/or STING-associated type I IFN signaling in PH development.

Spontaneous PH development in SAVI patients is thought to originate from systemic inflammation caused by augmentation of type I IFN signaling. However, studies on type I IFN in PH have been inconclusive, with contradicting results ^22,23^. Therefore, to better understand SAVI and STING^-/-^ mice pulmonary vascular changes, we examined the concept preliminarily through collecting whole lung protein from STING^-/-^ mice that were subjected to either bleomycin or chronic hypoxia for detection of IFN-related chemokines and cytokines. Intriguingly, there was no significant difference in IFNr, CXCL10, IL12 p40, and CXCL9 levels between lungs of control and experimental mice across models (**Fig. S2, B – E**). While serum level of IFNb was similar between WT and STING^-/-^ mice (**Fig. S2F**), IFNa level was below detection limit (data not shown). Taken together, we suspected that STING potentially contributes to PH through type I IFN signaling independent mechanisms.

### Type I IFN signaling plays minor role in PH development secondary to bleomycin and chronic hypoxia

To inspect the concept of STING regulating PH independent of type I IFN in more details, we induced PH in 8-10-week-old C57BL/6 (WT) mice with either bleomycin or chronic hypoxia exposure, while blocking type I IFN signaling using anti-IFNAR1 antibodies (aIFNAR1). In our hands, WT mice treated with bleomycin and aIFNAR1 concurrently displayed elevated RVSP comparable to isotype controls, with no accompanying change in RV remodeling (**Fig. 3, A and B**). In contrast, mice treated with aIFNAR1 under chronic hypoxia showed a decrease in RVSP relative to control mice, but still no change in RV remodeling (**Fig. 3, A and B**). Muscularized pulmonary vessel assessment demonstrated no significant difference between experimental groups in our chronic hypoxia model, however (**Fig. 3, C – E**). In addition, there was a significant increase in small and medium muscularized vessels of WT mice treated with both bleomycin and aIFNAR1, with a trend towards higher number of complete muscularized vessels (**Fig. 3, C – E**). Correlating with physiological data, MTC staining revealed no rescue of inflammation and fibrosis scarring in bleomycin-induced PH mice (**Fig. 3, F and G**). Interestingly, treatment with aIFNAR1 alone was sufficient to facilitate recruitment of CD11b^+^ cells into the lungs, specifically CD11b^+^Ly6C^hi^Ly6G^-^, but not CD11b^+^Ly6C^lo^Ly6G^+^ (**Fig. 3, H – J**, gating strategy shown in **Fig. 3K**). We noted the reversed trend in hypoxia-exposed mice with type I IFN blockade, where there was a sharp drop in the pathogenic CD11b^+^Ly6C^lo^Ly6G^+^ population, correlating with the physiological data, and consistent with our previous findings^29^. However, we found no change in pSTAT1 and pSTAT3 or other downstream signaling of type I IFN signaling, unlike what was observed in SAVI mice (**Fig. S4, A and B**). From this, we conclude that the observed phenotypic change is likely independent of type I IFN signaling.

**Figure 3.**
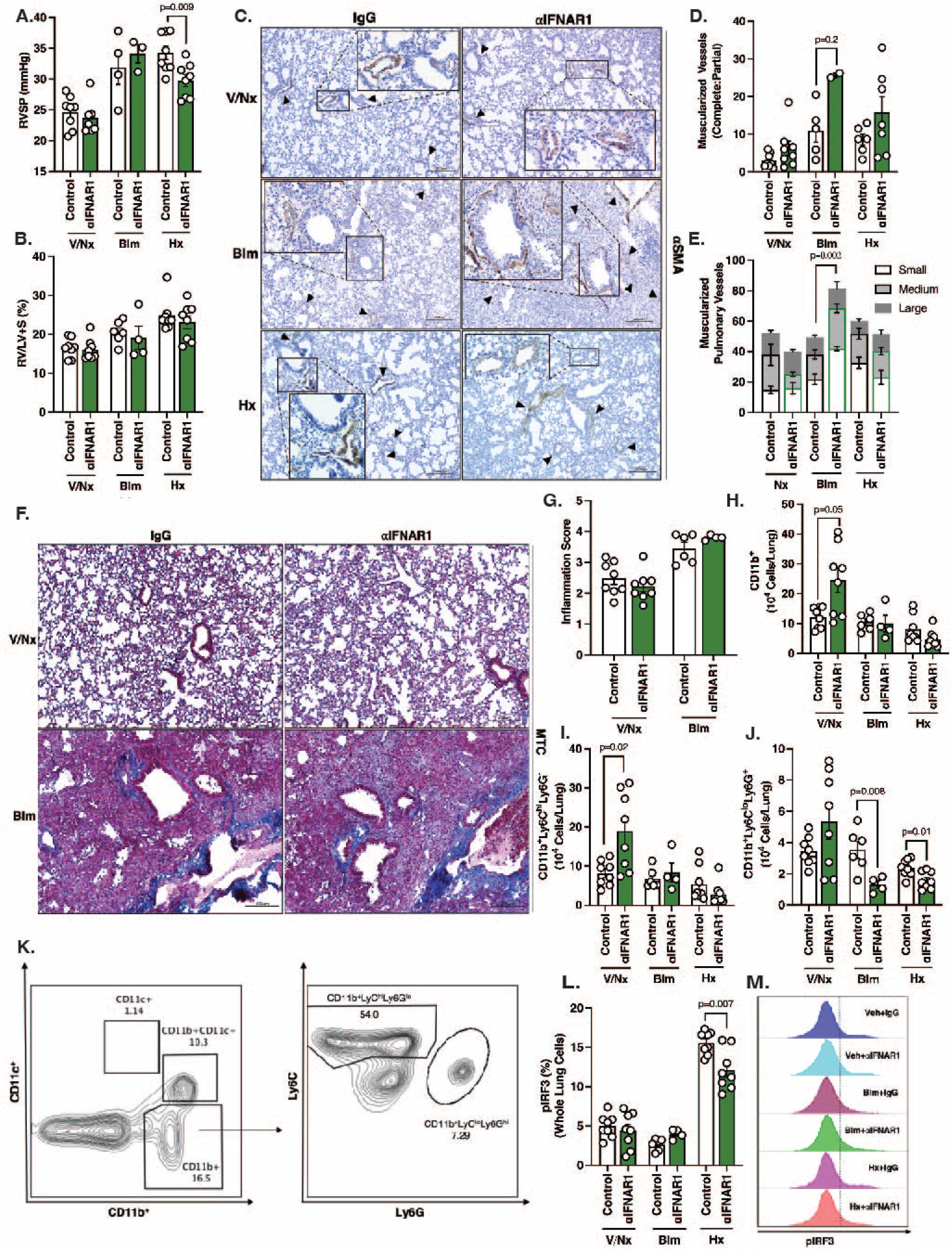
Type I IFN signaling plays minor role in PH development secondary to bleomycin and chronic hypoxia. (**A**) RVSP measurements of C57BL/6 mice receiving either control IgG or anti-IFNAR1 antibodies, while being subjected to PH with either bleomycin (Blm) or chronic hypoxia (Hx) exposure. (**B**) Fulton Index of C57BL/6 mice from dedicated mouse groups. (**C**) Representative images of IHC staining for a-SMA (brown, arrowheads) of formalin-fixed lung sections of 6 experimental mouse groups. Scale bar = 200mm at 10x magnification. (**D and E**) Quantification of small, medium, large, complete, or partial muscularized vessels from designated mouse groups. (**F**) Representative images of MTC-stained formalin-fixed lung sections of mice exposed to bleomycin, receiving either IgG or anti-IFNAR1 antibodies. Scale bar = 200mm at 10x magnification. (**G**) Inflammation/fibrosis scoring of lung sections obtained from MTC stains. (**H – J**) Flow cytometric quantification of (**H**) CD11b^+^, (**I**) CD11b^+^Ly6C^hi^L6G^-^, and (**J**) CD11b^+^Ly6^lo^Ly6G^+^ cells. (**K**) Flow cytometric gating strategy for identification of cell populations. (**L and M**) Quantification and flow plot representations of pIRF3 expression of whole lung cells from designated mouse groups. Each dot is an individual mouse (n=3-8/group). Column represents mean ± SEM. Significance level was calculated with un-paired two-tailed student T’s test. P<0.05 was considered significant.

To validate these results in an independent model, we utilized global Interferon a Receptor 1 deficient mice (IFNAR1^-/-^, 8-10-week-old), subjecting animals to bleomycin or hypoxia, as per the inhibitory antibody studies. There was no difference in RVSP between IFNAR1^-/-^ and WT mice in either of our PH models (**Fig. S4C**). In addition, there was no change in Fulton Index (**Fig. S4D**). We also noted a decrease, not an increase as per the antibody studies, in infiltrated CD11b^+^ and CD11b^+^Ly6C^lo^Ly6G^+^ cells in IFNAR1^-/-^ mice (**Fig. S4, E – G**). These data are consistent with current agreement in the field that inhibiting type I IFN signaling would be expected to decrease leukocyte recruitment in response to inflammatory stimuli^33,34^. Altogether, these data suggest that STING-associated type I IFN signaling plays trivial role in PH development secondary to bleomycin and chronic hypoxia. However, we did notice that chronic hypoxia-exposed WT mice treated with aIFNAR1 that were protected against RVSP elevation, experienced a profound decrease in phosphorylated IRF3, a direct downstream protein of STING in type I IFN signaling, suggesting a decrease in STING signaling (**Fig. 3, K and L**). Nonetheless, the phenotype of STING^-/-^ mice undergoing PH induction (both physiologically and immunologically) were highly suggestive that STING contributes to PH development largely independent of type I IFN signaling.

### scRNAseq shows most changes in stromal cells and myeloid cells in STING^-/-^ mice exposed to chronic hypoxia, independent of STING-associated type I IFN signature genes

PH has been primarily studied in the context of homeostatic loss in resident lung cells, namely pulmonary arterial endothelial cells (PAECs)^35,36^ and pulmonary vascular smooth muscle cells (PVSMCs)^37^. However, an influx of various immune cells from both the innate and adaptive immunity has also been described as a hallmark characteristic of PH^7,8^. Interestingly, STING expression and activation in both hematopoietic and non-hematopoietic cells have been shown to contribute to various disease pathogenesis that share the chronic inflammatory settings with similar biologic features to PH^18,38–40^. We therefore next hypothesized that STING contributes to PH in a cell-specific manner. To explore the cell-dependent role of STING in PH development, we generated three groups of bone marrow chimeric mice with either WT or STING^-/-^ hematopoietic cells, on a WT or STING^-/-^ non-hematopoietic background (**Fig. S5B**). Donor mice are either CD45.1^+^ C57BL/6J or STING^-/-^ mice, while recipient mice were either CD45.2^+^ C57BL/6J or STING^-/-^ mice that underwent whole body radiation. Chimerism was confirmed 6 weeks post bone marrow injection (**Fig. S5A**). Upon PH induction through chronic hypoxia exposure, a decrease in RVSP was observed in STING^-/-^ mice receiving WT bone marrow cells (i.e., WT hematopoietic cells and STING^-/-^ non-hematopoietic cells) (**Fig. S5C**), despite no observed change in RV remodeling (**Fig. S5D**). Staining with aSMA revealed no difference in muscularized pulmonary vessels, but STING^-/-^ mice receiving WT bone marrow cells (i.e., WT hematopoietic cells and STING^-/-^ non-hematopoietic cells) displayed a trend towards higher ratio of complete-to-partial vessel muscularization compared to WT mice receiving STING^-/-^ hematopoietic cells (i.e., STING^-/-^ hematopoietic cells and WT non-hematopoietic cells) (**Fig. S5, E – G**). Although there was not a significant difference in number of infiltrated CD11b^+^ cells, CD11b^+^Ly6C^hi^Ly6G^-^, and CD11b^+^Ly6C^lo^Ly6G^+^ correlating with RVSP changes, we again noted a trend towards higher infiltrated inflammatory cells in WT mice receiving STING^-/-^ bone marrow cells (i.e., STING^-/-^ hematopoietic cells and WT non-hematopoietic cells) (**Fig. S5, H – J**). We conclude the muted immunophenotype is likely due to the incomplete chimerism of bone marrow chimera generation, specifically the lower percentage in STING^-/-^ mice receiving WT bone marrow cells. The reason for this difference in engraftment efficiency is an area of ongoing research, regarding STING response to radiation-induced injury^41^. Given our physiologic data, we expected to see an increase in STING activation, assessed via increase in pIRF3 expression, in infiltrated inflammatory cells of mice with higher RVSP. However, we instead detected a significant reduction of pIRF3 in WT mice receiving STING^-/-^ bone marrow cells (i.e., STING^-/-^ hematopoietic cells and WT non-hematopoietic cells) compared to other groups (**Fig. S5, L and M**). From this, we hypothesized that as opposed to global STING knock-out, conditional knock-out of STING in certain hematopoietic cells could potentially worsen PH. Thus, from our chimera data, we developed two novel hypotheses for further testing; one, that STING contributes to PH in a non-hematopoietic dependent manner, and two, that STING expression and activation in hematopoietic cells provides protection against PH.

To further investigate the cell-specific role of STING in PH development, we performed single cell RNA sequencing (scRNAseq) on whole lung cells of 8-week-old WT and STING^-/-^ mice exposed to either normoxia or chronic hypoxia (n=2/group). Of particular note, we did not detect any differences in type I IFN signature genes among cell populations that mirror the phenotypic changes in STING^-/-^ mice (**Figure 4**). This finding further reinforces our data supporting an independent role of STING-associated type I IFN signaling in PH development and progression.

**Figure 4.**
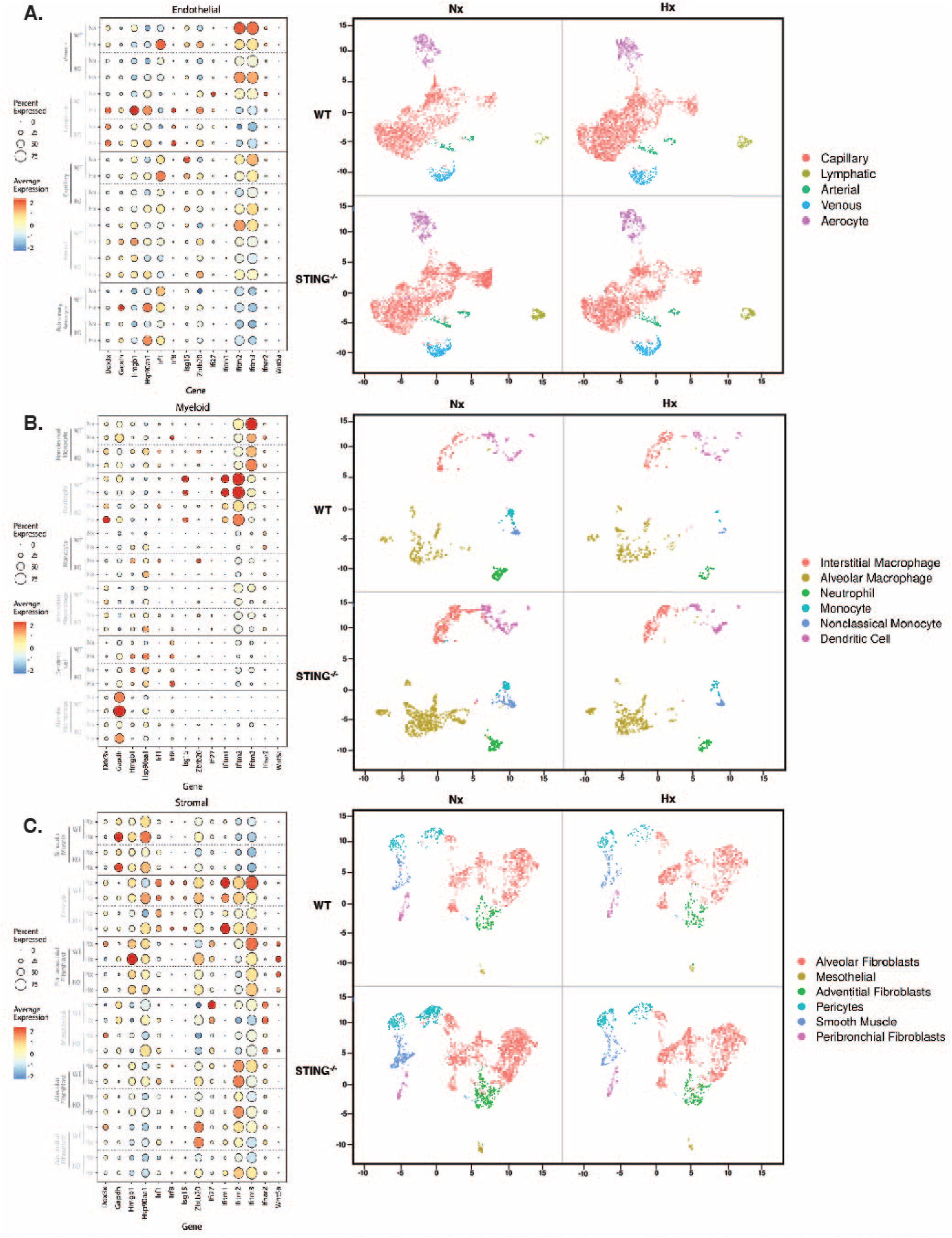
scRNAseq shows most changes in stromal cells and myeloid cells in STING^-/-^ mice exposed to chronic hypoxia, independent of STING-associated type I IFN signature genes. Expression of STING-associated type IF IFN genes and UMAP projections of (**A**) Endothelial cells, (**B**) Myeloid cells, and (**C**) Stromal cells of WT and STING^-/-^ exposed to either normoxia or chronic hypoxia (n=2/group).

In the non-hematopoietic cell compartment, a significant amount of PH research to date has focused on endothelial cells^35^ as well as stromal cells (i.e., PVSMCs)^37^. Thus, pathogenic roles for both cell types in PH development and progression have been identified. Interestingly, our scRNAseq data showed little change in endothelial cell populations of STING^-/-^ mice compared to WT mice undergoing chronic hypoxia exposure (**Fig. 4A**). Consistent with the recognition of PVSMC’s paramount role in PH, however, we were relatively unsurprised to note relevant differences in smooth muscle cell population between genotypes, as well as various other subpopulations of stromal cells between mouse groups (**Fig. 4C**). Additionally, in agreement with our bone marrow chimera data, that hinted at a potential protective role for STING expression in myeloid-derived cells, we detected an increase number of infiltrated myeloid cells, specifically nonclassical monocytes, and neutrophils, mirroring inflammatory resolution in chronic hypoxia-exposed STING^-/-^ mice that are protected from elevated RVSP, compared to WT (**Fig. 4B**). Thus, we identified smooth muscle cells (non-hematopoietic) and myeloid cells (hematopoietic) as our top candidates for next steps in studying the cellspecific role of STING in PH development.

### Smooth muscle, but not endothelial, cell specific deletion of STING protects against PH development

Encouraged by the scRNAseq data, we next proceeded to generate mice with cell-specific deletion of STING to evaluate the non-hematopoietic role of STING in PH development. VE-Cadherin-Cre (endothelial specific) and aSMA-cre (smooth muscle specific) mice were crossed to STING^fl/fl^ mice to produce mice with endothelial-specific and smooth muscle-specific deletion of STING (“eSTING” and “smSTING”, respectively). Surprisingly, yet consistent with the scRNAseq data, there was no difference in RVSP or Fulton Index between WT and eSTING subjected to either bleomycin or chronic hypoxia (**Fig. 5, A and B**). Likewise, there was also no change in the immune cell profile (i.e., CD11b^+^, CD11b^+^Ly6C^hi^Ly6G^-^, and CD11b^+^Ly6C^lo^Ly6G^+^) in the lungs of eSTING mice (**Fig. S6, A – C**). From here, given our scRNAseq and chimera data, as well as the consistency in phenotype of chronic hypoxia model, 8-10-week-old smSTING mice was subjected to chronic hypoxia for PH induction. Excitedly, protection against RVSP elevation upon chronic hypoxia exposure was observed in smSTING mice (**Fig. 7C**), as suggested in our scRNAseq and chimera data. There was a nonsignificant trend towards decreasing level of RV remodeling in smSTING exposed to chronic hypoxia (**Fig. 5D**). Mice with deletion of STING in SMCs did demonstrate a lower level of muscularized pulmonary vessels, especially small vessels, at baseline without a significant difference upon hypoxia exposure (**Fig. 5, E and G**), though they did have less complete muscularization compared to WT (**Fig. 5, E and F**). Somewhat surprisingly, there was a notable increase in infiltrated inflammatory cells (i.e., CD11b^+^, CD11b^+^Ly6C^hi^Ly6G^-^, and CD11b^+^Ly6C^lo^Ly6G^+^) in smSTING mice, despite the noted protection against RVSP elevation upon chronic hypoxia exposure, suggesting a role for SMC STING expression in inflammatory cell recruitment and activation (**Fig. 5, H – J**). From these data, we conclude a pathogenic role for smooth muscle, but not endothelial, STING in PH development.

**Figure 5.**
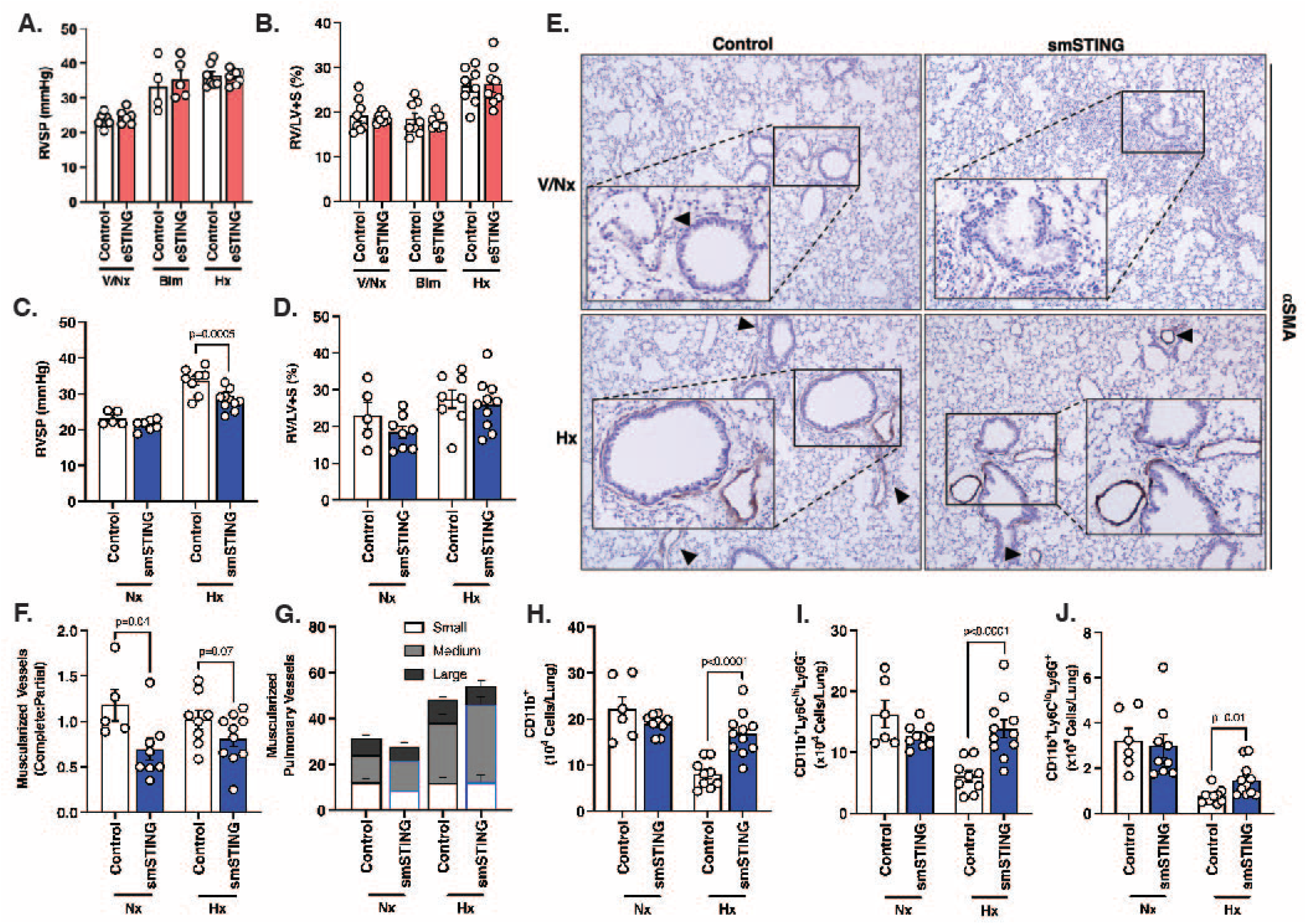
Smooth muscle, but not endothelial, specific deletion of STING provides protection against PH development secondary to chronic hypoxia exposure in mouse. (**A**) RVSP measurement of WT and endothelial-specific STING deficient (eSTING) mice at baseline, treated with bleomycin (Blm) or exposed to chronic hypoxia (Hx). (**B**) Fulton Index of WT and eSTING mice from all experimental groups. (**C**) RVSP measurement of WT and smooth muscle-specific STING deficient (smSTING) mice at baseline or exposed to chronic hypoxia. (**D**) Fulton Index of WT and smSTING mice from 6 treatment groups. (**E**) Representative images of aSMA (brown, arrowheads) IHC staining of formalin-fixed lung sections from WT and smSTING mice. (**F and G**) Quantification of small, medium, large, complete and partial muscularized pulmonary vessels from aSMA-stained lung sections of WT and smSTING mice across dedicated mouse groups. (**H** – **J**) Flow cytometric quantification of pulmonary infiltrated (**H**) CD11b^+^, (**I**) CD11b^+^Ly6C^hi^L6G^-^, and (**K**) CD11b^+^Ly6C^lo^Ly6G^+^ cells of WT and smSTING mice subjected to normoxia, bleomycin, or chronic hypoxia. Each dot is an individual mouse (n=5-8/group). Column represents mean ± SEM. Significance level was calculated with un-paired two-tailed student T’s test. P<0.05 was considered significant.

**Figure 6.**
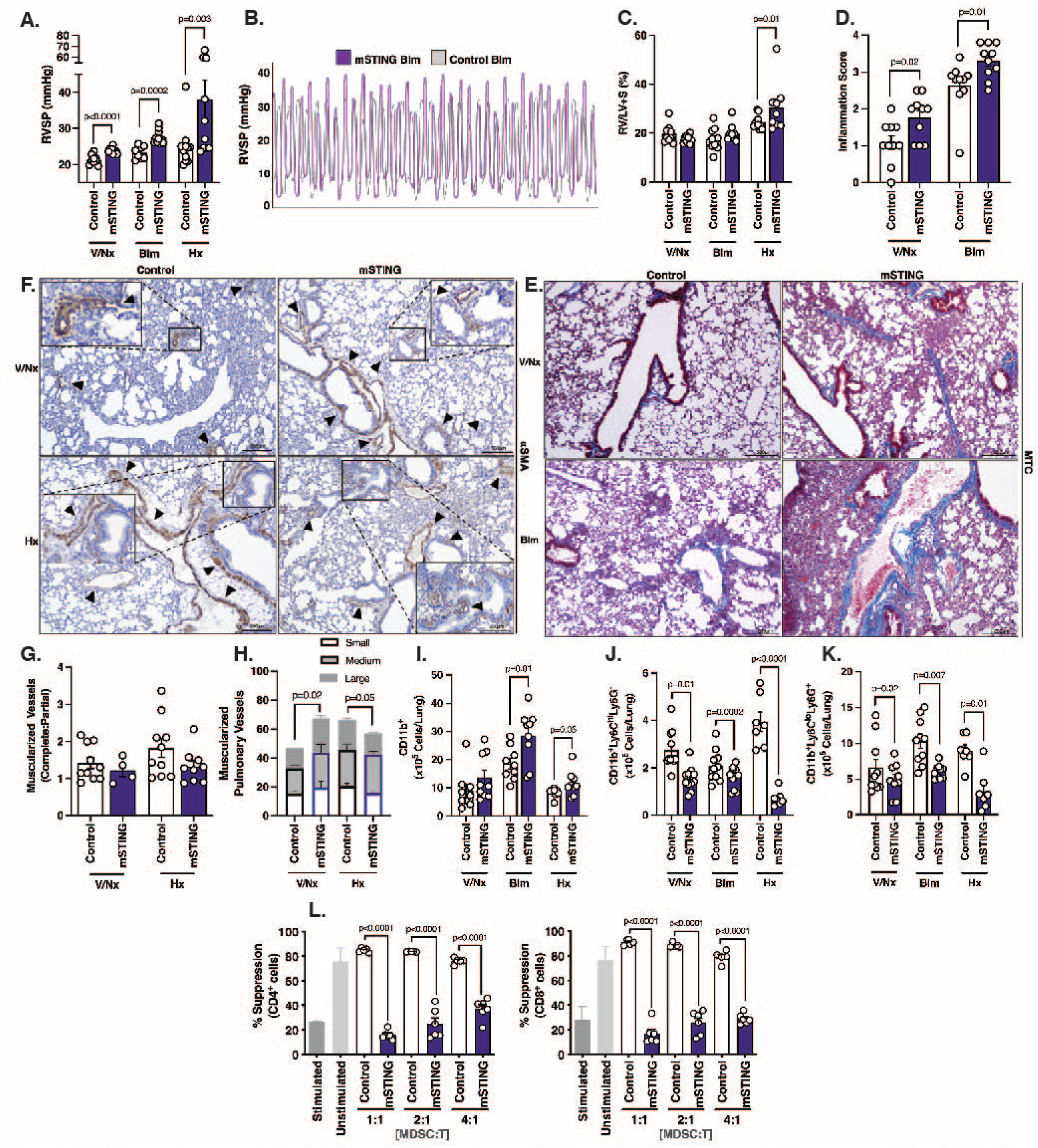
STING expression on myeloid cell is necessary to prevent severe PH. (**A and B**) RVSP measurement and curve representation of WT and myeloid-specific deficient STING (mSTING) mice subjected to normoxia, bleomycin, or chronic hypoxia. (**C**) Fulton Index of WT and mSTING mice from designated groups. (**D**) Inflammation/fibrosis quantification assessed through MTC-stained lung sections of WT and mSTING mice with or without bleomycin treatment. (**E**) Representative images of MTC-stained formalin-fixed lung sections of WT and mSTING mice from dedicated groups at 10x magnification. (**F**) Representation of aSMA (brown, arrowheads) IHC staining of formalin-fixed lung sections from WT and mSTING mice at baseline and exposed to chronic hypoxia at 10x magnification. (**G and H**) Quantification of muscularized pulmonary vessels (small, medium, large, complete, partial) of WT and mSTING mice of designated group, using aSMA IHC staining. (**I** – **K**) Flow cytometric quantification of pulmonary infiltrated (**I**) CD11b^+^, (**J**) CD11b^+^Ly6C^hi^L6G^-^, and (**K**) CD11b^+^Ly6C^lo^Ly6G^+^ cells of WT and smSTING mice subjected to normoxia, bleomycin, or chronic hypoxia. (**L**) Quantification of suppression capability of splenic MDSC isolated from control and mSTING mice on CD4^+^ and CD8^+^ T cells. % Suppression is measured by proliferation of T cells labeled with Cell Trace Violet (CTV) and stimulated with anti-CD3/CD28 antibodies at listed ratios. Each dot is an individual mouse (n=5-10/group). Column represents mean ± SEM. Significance level was calculated with un-paired two-tailed student T’s test. P<0.05 was considered significant.

**Figure 7.**
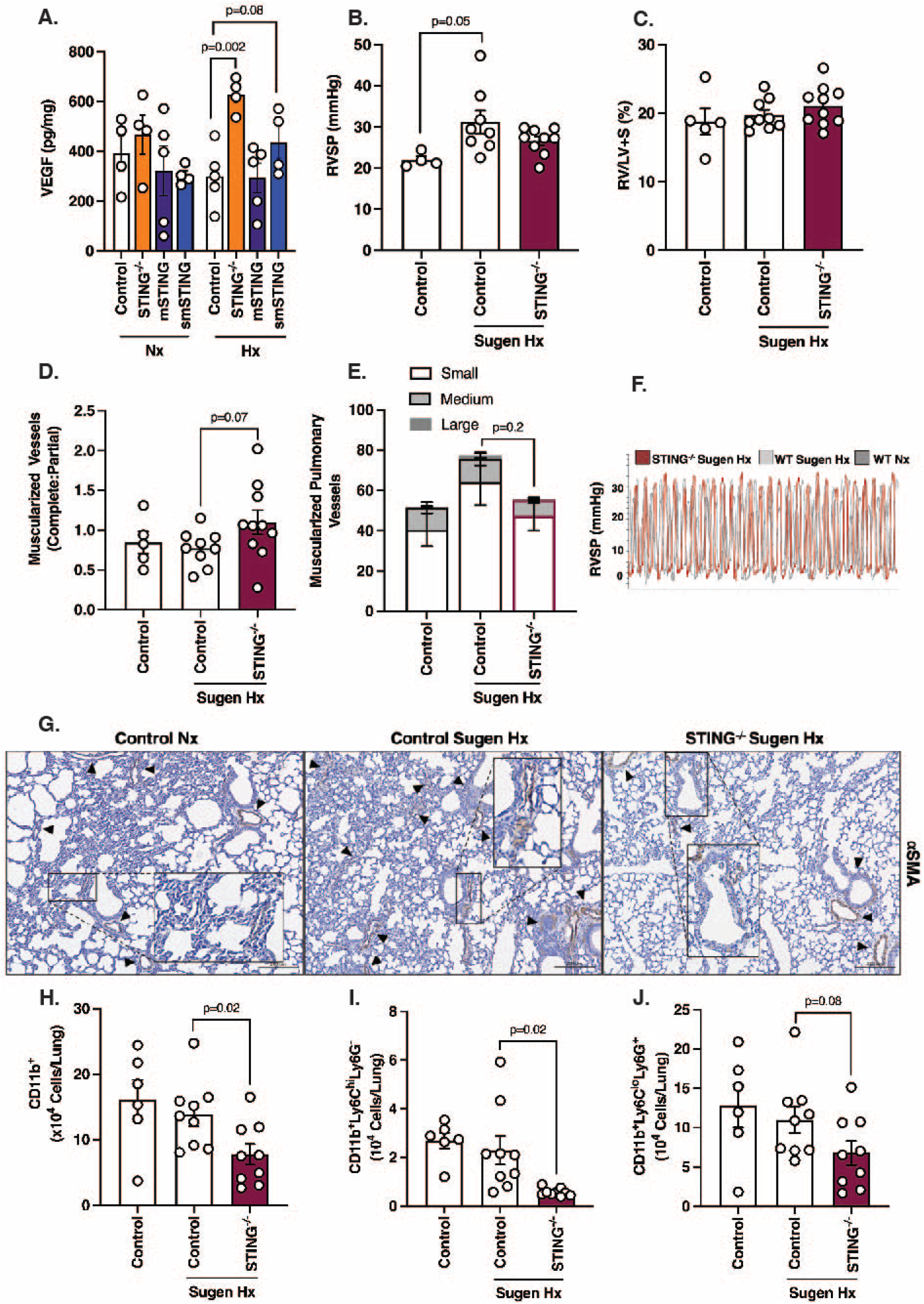
Inhibition of VEGF signaling reverses protection against PH secondary to chronic hypoxia in STING^-/-^ mice. (**A**) Quantification of vascular endothelial growth factor (VEGF) in whole lung cells of WT, STING^-/-^, mSTING and smSTING mice at baseline and when subjected to chronic hypoxia (Hx). Each dot represents an individual mouse (n=4/group). (**B and F**) RVSP measurement and cure representation of WT and STING^-/-^ mice subjected to normoxia (Nx) or chronic hypoxia with Sugen, VEGF inhibitor (Sugen Hx). (**C**) Fulton Index of WT and STING^-/-^ mice from dedicated mouse groups. (**D and E**) Quantification of muscularized pulmonary vessels (small, medium, large, complete and partial) of WT and STING^-/-^ mice in designated group, assessed with aSMA IHC staining of formalin-fixed lung sections. (**G**) Representative images of aSMA (brown, arrowheads) IHC staining of formalin-fixed lung sections of WT and STING^-/-^ mice in designated groups. (**H – J**) Flow cytometric quantification of pulmonary infiltrated (**H**) CD11b^+^, (**I**) CD11b^+^Ly6C^hi^L6G^-^, and (**J**) CD11b^+^Ly6C^lo^Ly6G^+^ cells of WT and STING^-/-^ mice subjected to normoxia or Sugen chronic hypoxia. Each dot is an individual mouse (n=6-8/group). Column represents mean ± SEM. Significance level was calculated with un-paired two-tailed student T’s test. P<0.05 was considered significant.

### STING expression on myeloid cell is necessary to prevent severe PH

Having identified the pathogenic role of smooth muscle STING in PH, we next sought to understand the role of STING expression by myeloid cell, in PH development. LysM-Cre mice were crossed with STING^fl/fl^ mice (“mSTING”) to generate mice myeloid-specific deletion of STING. Given our lab’s prior work with the LysM-Cre-modeling in both bleomycin-induced and chronic hypoxia-induced PH^29–32^ showing that MDSC suppressive phenotype is strongest under bleomycin treatment, 8-10-week-old mSTING mice were subjected to PH using both models. Consistent with our prediction that hematopoietic-lineage STING expression may have a protective role against PH, mSTING mice developed severe PH under both bleomycin treatment and chronic hypoxia exposure, defined by elevation of RVSP (**Fig. 6, A and B**) and RV remodeling with increased in Fulton Index in mSTING mice exposed to chronic hypoxia (**Fig. 6C**). It is worth highlighting that even at baseline, mSTING mice developed higher RVSP compared to control mice, signifying the important role of myeloid STING in pulmonary vascular homeostasis (**Fig. 6A**). Related, an increase in inflammation/fibrosis was observed in mSTING mice treated with bleomycin comparable to WT (**Fig. 6, D and E**). Consistent with RVSP data, we saw a significant higher degree of tissue scarring and collagen deposition in mSTING mice even at baseline (**Fig. 6, D and E**). Assessment of muscularized pulmonary vessels demonstrated an increase in medium and large vessels in mSTING mice at baseline (**Fig. 6, F and H**). However, upon chronic hypoxia exposure, there were fewer small vessels present in mSTING mice compared to WT, while number of medium and large vessels remained similar (**Fig. 6, F and H**). On the same note, less complete vessel muscularization was noted in mSTING mouse lungs (**Fig. 6, F and G**). We therefore cannot rule out a contribution from parenchymal fibrosis in development of pulmonary vascular remodeling, and vessel rarefaction, as the increase in scarring likely contributed to the phenotype. Upon flow cytometric analysis of lungs, we noted an increase in CD11b^+^ cells in mSTING mice subjected to either bleomycin or chronic hypoxia compared to WT (**Fig. 6I**). Interestingly, however, there was a decrease in number of CD11b^+^Ly6C^hi^Ly6G^-^ and CD11b^+^Ly6C^lo^Ly6G^+^ in mSTING mice with severe PH, similar to that of global STING knockout mice (**Fig. 6, J and K**). We therefore hypothesized that there was a unique immunophenotypic functional difference in myeloid cells from the mSTING mice. We tested this hypothesis by assessing the regulatory capability of isolated myeloid cells compared to their WT counterparts, finding a decrease in the immunosuppressive capacity as assessed by T cell suppression assay (**Fig. 6L**). Thus, the PH phenotype in our respective cell-specific deletion mouse models, is potentially contributed to by suppressive cell function, in a STING-dependent manner. In conclusion, we have demonstrated that myeloid STING is an important intermediate in prevention of severe PH, associated with changes in the suppressive signature of these cells.

### Inhibition of VEGF signaling reverses protection against PH secondary to chronic hypoxia in STING^-/-^ mice

Having identified smooth muscle cell and myeloid cell specific STING expression as a regulatory component involved in PH development, we aimed next to find a unifying underlying mechanism to which STING aggravates or prevents PH. Correlating with changes in RVSP, we found significant alterations of vascular endothelial growth factor (VEGF) in STING^-/-^, mSTING, and smSTING mice subjected to chronic hypoxia (**Fig. 7A**). Mice that were protected from elevated RVSP demonstrated an increase in level of VEGF, with STING^-/-^ mice showing the highest level of expression. Similarly, mice with increased RVSP displayed a decrease in VEGF expression, namely mSTING mice (**Fig. 7A**). We therefore hypothesized that interruption of VEGF signaling in these mice would nullify the phenotype, and globally deficient mice would display worsened PH.

To further investigate this finding, we employed the Sugen/Hypoxia PH model on 8-10-week-old STING^-/-^ mice, in which the mice received weekly injection of Sugen (SU5416), a Vascular Endothelial Growth Factor Receptor 1 and 2 (VEGFR1/2) antagonist, before and while undergoing chronic hypoxia exposure. The combination of Sugen and chronic hypoxia has been shown to cause pronounced PH in both rat and mice, and to be considered the “gold standard” model by many in the PH field, due to the irreversible nature of pulmonary vascular remodeling and presence of plexiform lesions on histology^42^. As hypothesized, Sugen/Hypoxia exposed STING^-/-^ mice no longer demonstrated protection against PH, as there was no significant difference between WT and STING^-/-^ mice (**Fig. 7, B and F**). Similarly, there was no change in RV remodeling between experimental groups (**Fig. 7C**). Assessment of muscularized pulmonary vessels did demonstrate a trend towards attenuation of small vessel muscularization in Sugen/Hypoxia STING^-/-^ mice compared to WT, and a slight increase in complete muscularized vessels (**Fig. 7, D, E and G**). Interestingly, in correlation with VEGF immunosuppressive ability, we observed an overall decrease, or a trend towards reduction, in lung CD11b^+^, CD11b^+^Ly6C^hi^Ly6G^-^, and CD11b^+^Ly6C^lo^Ly6G^+^ cells between STING^-/-^ and WT mice undergoing Sugen chronic hypoxia, despite there being no change in RVSP (**Figure 7, H – J**). These data highlight the complex role of VEGF signaling in PH, as well as the importance of potential functional differences in myeloid cells, over simple quantity, in disease pathogenesis. These experiments thus shed novel light on STING-VEGF signaling axis in an established PH disease model.

### Severity of PH in mSTING mice is mediated by PD-L1

Given that myeloid-specific deletion of STING caused severe PH in mice, though seemingly independent of myeloid cell infiltration if not function, we asked if there was a separate STING-associated mechanism that is specific to myeloid cells, especially MDSCs, that could explain the worsening in pulmonary vascular remodeling. We have previously demonstrated that MDSC in PH patients display a markedly increased level of PD-L1 expression, correlating with disease severity, a result that we later were able to phenocopy in mice, with higher PD-L1 expression on MDSCs reflecting an increase in suppressive capability and worsened PH^32^. Therefore, we set out to examine PD-L1 expression on CD11b^+^Ly6C^hi^Ly6G^lo^ and CD11b^+^Ly6C^lo^Ly6G^+^ cells in STING^-/-^ mice and other cell-specific deletion mouse models. Upon analyzing PD-L1 expression in PH-induced STING^-/-^ mice, we noted a trend towards decreasing PD-L1 expression in CD11b^+^, CD11b^+^Ly6C^hi^Ly6G^-^ as well as CD11b^+^Ly6C^lo^Ly6G^+^ cells (**Fig. 8, A – D**). As previously mentioned, CD11b^+^Ly6C^lo^Ly6G^+^ cells have been considered to be the pathogenic cell-type in PH^29^. In chronic hypoxia-exposed mice, CD11b^+^Ly6C^lo^Ly6G^+^ cells of STING deficient mice showed the most significant reduction in PD-L1 expression compared to controls (**Fig. 8, C and D**). As mSTING mice developed severe PH, we hypothesized they would display higher expression of PD-L1 on this cell population of interest, and as predicted, these mice demonstrated a significant increase in PD-L1 expression on both CD11b^+^Ly6C^hi^Ly6G^-^ and CD11b^+^Ly6C^lo^Ly6G^+^ cells after induction of PH (**Fig. 8, E – H**). Notably, there was an increase in PD-L1 expression on lung myeloid cells at baseline in mSTING mice, correlating with the baseline elevation of RVSP in these mice (**Fig. 8, E – H**). STING-regulated PD-L1 expression on MDSCs seems to be unique to myeloid cell-specific model, as smSTING mice expressed very little PD-L1 on their infiltrated myeloid cells, although there was some difference among experimental groups (**Fig. S7, A – C**). Relevantly, an increase in PD-L1 expression was also observed in these two cell populations within the lungs of SAVI mice (**Fig. S7, D and E**). Thus, in summary, STING-regulated PD-L1 expression has the potential to serve as a pathogenic marker, and pharmaceutical target, on myeloid cells in PH, especially the pathogenic CD11b^+^Ly6C^lo^Ly6G^+^ cells, correlating well with disease severity. We therefore next sought to uncover a role for PD-L1 inhibitors in mSTING mice, as a method of abrogating severe PH.

**Figure 8.**
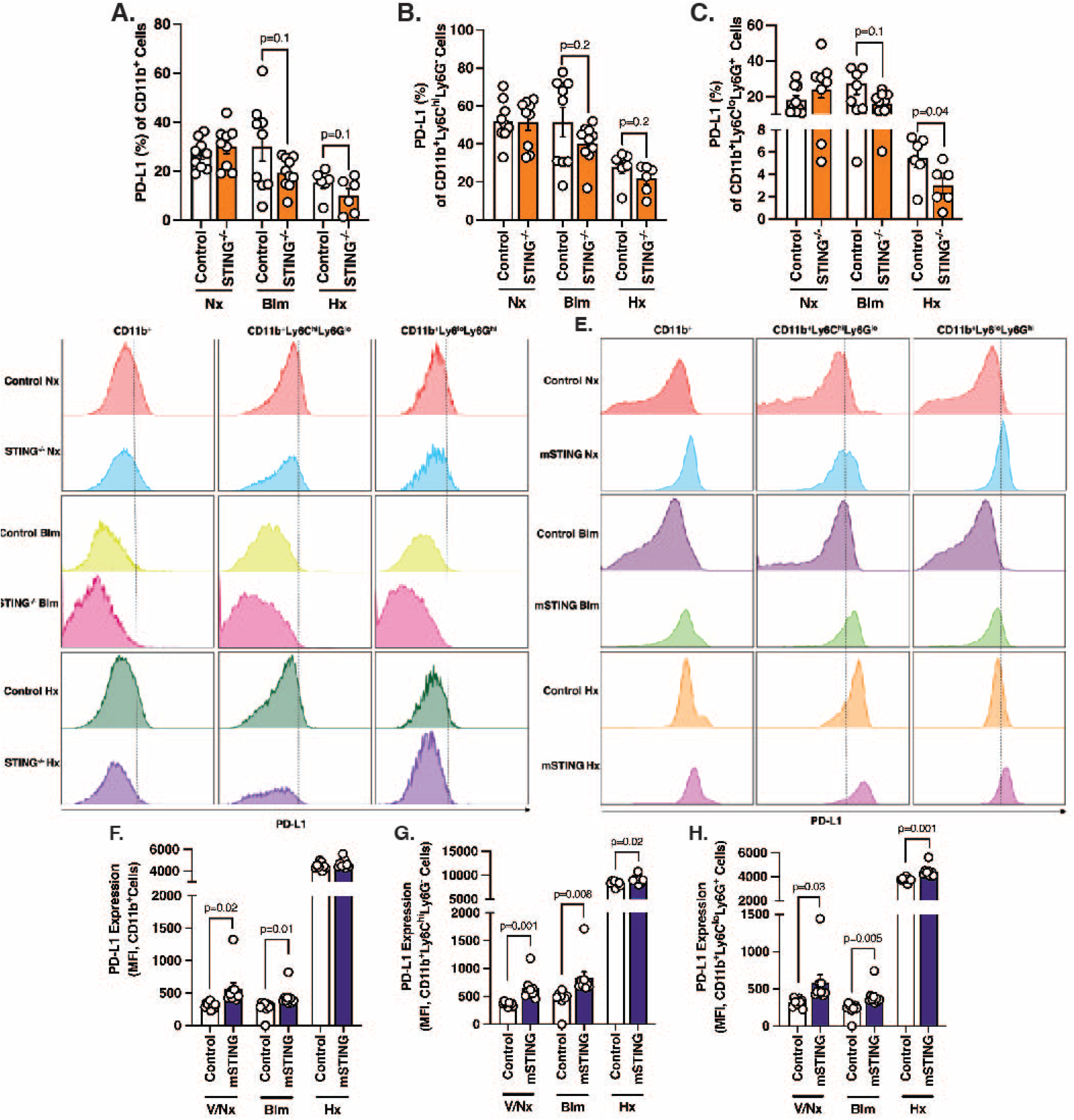
Severity of PH in mSTING mice is PD-L1 dependent. (**A** – **D**) Flow cytometric quantification and flow charts representations of PD-L1 expression in pulmonary infiltrated (**A**) CD11b^+^, (**B**) CD11b^+^Ly6C^hi^L6G^-^, and (**C**) CD11b^+^Ly6C^lo^Ly6G^+^ cells of WT and STING^-/-^ mice in dedicated mouse groups. (**E – F**) Representative flow charts and quantification of PD-L1 expression in pulmonary infiltrated (**E**) CD11b^+^, (**G**) CD11b^+^Ly6C^hi^L6G^-^, and (**F**) CD11b^+^Ly6C^lo^Ly6G^+^ cells of WT and mSTING mice in designated mouse groups. Each dot represents an individual mouse (n=5-10/group). Column represents mean ± SEM. Significance level was calculated with un-paired two-tailed student T’s test. P<0.05 was considered significant.

### Anti-PD-L1 antibody abrogates severe PH in mSTING mice

We next treated mice – in a preventive and proof-of-concept manner – with a PD-L1 specific inhibitory antibody. Consistent with our prior findings using a different less-specific model of disease^32^, treatment with aPD-L1 in 8-10-week-old bleomycin-induced PH mice attenuated severe PH, demonstrating rescue of elevated RVSP, but not RV remodeling (**Fig. 9, A – C**). Interestingly, bleomycin-treated mSTING mice under aPD-L1 therapy showed a significant reduction of inflammation/fibrosis compared to WT, as well (**Fig. 9, D and E**). Reflecting these physiological data, we observed normalization in quantity of inflammatory cells (i.e., CD11b^+^, CD11b^+^Ly6C^hi^Ly6G^-^ and CD11b^+^Ly6C^lo^Ly6G^+^ cells) in bleomycin-treated mSTING mice receiving aPD-L1 therapy (**Fig. 9, F – H**). PD-L1 expression on myeloid cells was also normalized (**Fig. 9, I – K**). In conclusion, as mice with myeloid-specific STING deficient developed severe PH under bleomycin treatment, aPD-L1 therapy generally improved disease outcome in in these mice. Since PD-L1 expression correlates with disease severity in a cell-specific manner (i.e., myeloid cell), as well as in our global STING deficiency mouse model, a STING-PD-L1 signaling axis showcases the potential for therapeutic intervention in PH. Given the large amount of anti-PD-L1 therapeutics available, future studies will allow better assessment of this therapy in patients with PH.

**Figure 9.**
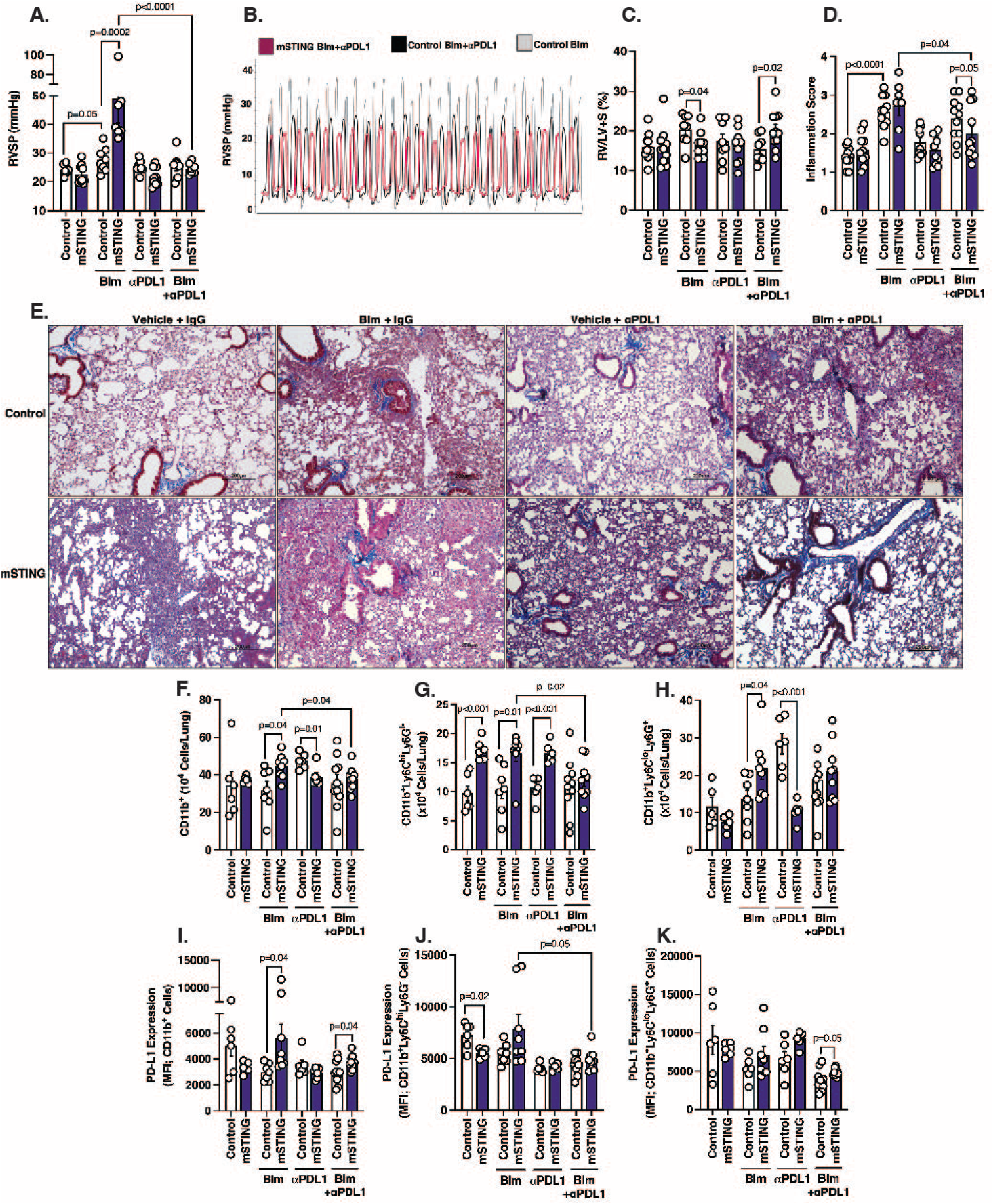
Anti-PD-L1 antibody abrogates severe PH in mSTING mice. (**A and B**) RVSP measurement and curve representation of WT and mSTING mice treated with bleomycin, with or without anti-PD-L1 antibody (aPD-L1) as indicated. (**C**) Fulton Index of WT and mSTING mice in dedicated treatment group. (**D and E**) Quantification of inflammation/fibrosis degree and representative images of MTC-stained formalin-fixed lung sections of WT and mSTING mice in designated experimental groups. (**F** – **H**) Flow cytometric quantification of pulmonary infiltrated (**F**) CD11b^+^, (**G**) CD11b^+^Ly6C^hi^L6G^-^, and (**H**) CD11b^+^Ly6C^lo^Ly6G^+^ cells of WT and mSTING mice in all experimental groups. (**I** – **K**) Flow cytometric quantification of PD-L1 expression on (**I**) CD11b^+^, (**J**) CD11b^+^Ly6C^hi^L6G^-^, and (**K**) CD11b^+^Ly6C^lo^Ly6G^+^ cells of WT and mSTING mice in all dedicated groups. Each dot is an individual mouse (n=6-12/group, data pooled from 2 independent experiments). Each dot is an individual mouse (n=6-8/group). Column represents mean ± SEM. Significance level was calculated with un-paired two-tailed student T’s test. P<0.05 was considered significant.

## Discussion

Pulmonary hypertension is a major global health issue, with Group 3 PH prevalence on the rise affecting up to 60% of patients with chronic lung diseases^2,3^. Currently there are no disease-modifying therapies for PH patients. Thus, there is an urgent need to identify novel disease mechanisms that can be therapeutically targeted. Here, we report a role for viral-DNA sensing protein STING in PH development. Using multiple PH models, we confirmed that STING activation in a cell-specific manner contributes to RVSP elevation, RV remodeling and immune cell infiltration. Moreover, we provide evidence that smooth muscle cell expression of STING worsens PH, while STING expression by myeloid cells is critical to prevent severe PH. Furthermore, we convincingly demonstrate that type I IFN signaling plays a relatively minor role in PH pathogenesis, with a larger contribution derived from VEGF signaling in a STING-dependent manner. Lastly, we show that STING-induced PD-L1 expression on infiltrated myeloid cells has the potential to serve as a marker for disease severity, and therapeutic target. Overall, our results identify STING as an important regulator in PH development and progression, and thus offer badly needed disease-modifying targeted pathways.

Due to the complex nature of PH, no single signaling pathway has accounted for all of disease characteristics. However, random genetic “experiments” by nature, such as mutations accounting for SAVI patients, may potentially inform disease pathology. Of interest, the SAVI mouse model with N153S mutation displayed an IRF3-independent mechanism regarding STING-associated vasculopathy^43^, while SCID phenotype in SAVI mice with V154 mutation occurs in the absence of IFN signaling^44^. Similarly, a recent study supports an important immunoregulatory function of STING in lung fibrosis, a common risk factor for PH^45^. The latter report revealed that the protective role of STING is independent of type I IFN but associated with dysregulated neutrophils, with similar finding in PBMC of IPF patients^46^. This observation was consistent with our own findings demonstrating a minor role of STING-associated type I IFN signaling in PH. A possible explanation is that the importance of type I IFN signaling is more profound in the early stage of disease development, in which inflammation resolution relies more heavily on innate immune response. With disease progression and chronic inflammation established, persistent IFN signaling may fuel pathogenic inflammation. To that end, early evaluation of PH in response to hypoxia in IFNAR1^-/-^ mice presented an opposite phenotype to our chronic hypoxia model^22^. Interestingly, it has been shown that type I IFN suppresses CXCR2-expressed neutrophil recruitment through inhibiting CXCL2 secretion^47,48^. Our data support this prior observation, with mice receiving anti-IFNAR1 antibody under both chronic hypoxia and bleomycin treatment displayed a decrease in infiltrated CD11b^+^Ly6C^lo^Ly6G^hi^ (i.e., murine PMN-MDSC). Related, our previous work has demonstrated a consistent role for CXCR2 in regulating MDSC recruitment and activation^29^.

Not only does STING play an important role in a variety of lung diseases as discussed above, but it also proves critical in other diseases classically associated with chronic inflammation. For instance, in cancer, a constellation of pathologies sharing multiples cellular mechanisms to PH^49^, STING has been shown to amplify MDSC recruitment to tumor site post-radiation, contributing to radiation-resistance^50^. In addition, STING expressed in tumor cells contribute to anti-tumor immunity^51^, highlighting the pivotal role for STING in suppressing immune response. Related, STING^-/-^ mice are rescued from disease in several inflammatory models such as silica-induced fibrosis and sepsis^52,53^, as well as autoimmune diseases like lupus or rheumatoid arthritis^54,55^. Our study provides the first evidence of STING regulating PH, in which we illustrated protection against PH development in STING^-/-^ mice in two different models. It is worth mentioning that rescue in right ventricular remodeling, lower level of fibrosis and muscularized vessels were observed in STING^-/-^ mice exposed to both bleomycin and chronic hypoxia, highlighting the relevance of STING in cells that take part in vessel muscularization and remodeling, such as myofibroblasts. Future studies regarding STING should thus focus on the specific mechanism preventing right-heart failure in end-stage PH patients, in light of our finding and other studies showing that STING^-/-^ mice were protected from cardiovascular diseases^56,57^.

STING expression and function is altered depending upon the cell-type under study, therefore it is not surprising that STING plays a cell specific role in PH. Given the literature-to-date on endothelial STING expression, however, it was unexpected that our endothelial deletion model did not indicate a significant role for the protein in disease onset and progression. As well, it was interesting that activation of STING in smooth muscle cells, versus in myeloid cells was protective against severe PH. Related, the immune cell profile of smSTING mice was opposed to what was observed in STING^-/-^ mice, despite protection against PH development in both models. One possible explanation for this is that the smSTING mice have alterations in the inflammatory resolution phase, due to cell-cell interactions, as it known that STING signaling in one cell can activate STING response in bystander cells^58^. It is interesting to speculate then that STING expression on SMC is crucial in both recruitment and activation of inflammatory cells (i.e., myeloid cells), regulating pulmonary vascular remodeling. The study of such cell-cell interactions in PH, either through paracrine signaling or as a field effect, is a subject of on-going investigation in the field^43,59,60^. Viral DNA sensing pathways could thus be a key mediator of these responses.

PD-L1 has previously been identified as an important regulator for MDSC immunosuppressive function^61–63^. Interestingly, in our study, an increase in PD-L1 expression in infiltrated myeloid cells correlated with decreased suppressive function, albeit in the setting of more severe disease. Additionally, blockage of PD-L1 with antibodies rescued mSTING mice from severe PH. First, it is possible that cell-specific deletion of STING in myeloid cells alters myeloid cell function, independent of quantity of infiltrated cells. These data also suggest that negative sequelae of PD-L1 expression on MDSC are potentially independent of its suppressive capability. For example, a recent paper illustrated a negative regulatory role of nuclear PD-L1 on STING activation in cancer cell that fuels cancer cell senescence^64^. This highlights an intriguing need to distinguish the role of nuclear and surface PD-L1 in disease context. This might be relevant to interpretation of our findings, as we focused exclusively on PD-L1 surface expression. The exact mechanism of STING in myeloid cell function, independent of PD-L1, is a subject for future study.

VEGF role in PH is complex, depending upon timing of expression, responsive cell-type, and concentration^65^. As such, treatment with VEGFR1/2 inhibitor, SU5416/Sugen, concurrently with chronic hypoxia exposure produces irreversible PH in rodents^66^. Therefore, it was quite surprising that the use of Sugen/hypoxia in our STING^-/-^ mouse overrode the protective phenotype, providing evidence for the importance of VEGF-STING signaling in vascular regeneration. Related, global knock-out and smooth muscle specific deletion of STING promoted increased VEGF expression compared to control and mSTING mice, correlating with protection against PH. This finding suggests a healthy angiogenic role for cell-specific VEGF that is STING-regulated. Future studies on VEGF-STING pathway in PH, as well as the source of VEGF secretion, will likely advance toward directed therapeutics.

Our study has several key limitations. Engraftment efficiency in our chimera experiment was directly influenced by the protein of interest, STING, in particular the response to external radiation. As such, parabiosis experiments may be an alternative in the future, though plagued by antibody-mediated rejection and undesired stem cell stimulation. Readily available therapeutic with PD-L1 inhibitor can cause pneumonitis, however bleomycin-induced lung injury in mSTING mice actually improved after PD-L1 antibody treatment. The mechanism to which it rescues PH thus needs further investigation before translation of effective therapies^67^. Moreover, there were sex-related differences observed in SAVI mice. These were not demonstrated in STING^-/-^ and cell-specific STING deficient mice. This may be due to estrogen metabolism in DNA sensing or hormonal effect on receptor expression related to STING signaling in SAVI mice, with constitutive activation of the STING-homodimer intracellularly, with DNA related damage and response in a sex-specific manner an ongoing area of research in the field^68,69^. Lastly, in our hands the source of STING activation is not fully explored, though cell free DNA has been described as relevant to PH in several human-cell and animal-based studies^24,26^. Further investigations are required to fully uncover the respective contribution of either native or viral DNA to STING activation in PH.

In summary, our study convincingly demonstrates the contribution of STING to PH development, in large part through inhibition of VEGF secretion. We have also shown a dichotomous role for STING in lung resident and infiltrated inflammatory cells in PH. Thus, cell-specific targeting of these pathways may be of therapeutic value in the future.

## Materials and methods

### Mice

SAVI, STING^-/-^ (MPYS^-/-^) and IFNAR1^-/-^ mice were purchased from Jackson Laboratory then crossed with in-house C75BL/6J background for maintenance. All transgenic mice generated in this study were C57BL/6J background. Transgenic mice expressing Cre-recombinase under the control of VE-cadherin promoter (VeCad-Cre) were crossed with STING mice flanked by two loxP sites (STING^fl/fl^) to generate Cre-mediated specific deletion of STING gene in endothelial cells. Similarly, transgenic mice expressing Cre-combinase under the control of LysM promoter (LysM-Cre) or SMA promoter (SMA-Cre) were crossed with STING^fl/fl^ mice for generation of Cre-mediated specific deletion of STING gene in myeloid cells, and smooth muscle cell, respectively. Breeding were set up, such that STING construct was kept in homozygous state, while VeCad-Cre, LysM-Cre and SMA-Cre were maintained in heterozygous state, yielding mice with Cre^+^ mice with respective cell deletion. Mice were bred and housed in specific pathogen-free conditions. Inhouse C57BL/6J mice were used as control for global knock-out and SAVI mice, while Cre^-^ mice were used as control for experiments involving transgenic mice. Gender matched 8-10 weeks mice were used, after confirmation of appropriate genotype with genotyping for all experiments unless specify otherwise. Animal experiments and maintenance were approved by the Institutional Animal Care and Use Committee of University of Florida (IACUC; Protocol 08702). Animal studies are reported in compliance with the ARRIVE guidelines (Kilkenny, Browne, Cuthill, Emerson, & Altman, 2010) and with recommendations made by the Journal of Experimental Medicine.

### Mouse models of pulmonary hypertension

Pulmonary hypertension (PH) was induced in mice either with bleomycin injection or chronic hypoxia exposure.

#### Bleomycin

Mice received intraperitoneal (*i.p*.) injections of bleomycin (Millipore Sigma #9041934) at 0.018U/g twice a week for 4 weeks. Weight of animals were monitored throughout injection period. A 20% loss of body weight resulted in temporary termination of bleomycin treatment. Injection was resumed when the animal regained at least 10% of loss weight. Euthanasia and data collection were performed 5 days after final injection, on Day 33 of bleomycin injection protocol.

#### Chronic hypoxia

Mice undergone chronic hypoxia exposure were placed in a normobaric ventilated chamber, in which the level of O_2_ is controlled through flow of N_2_ (ProO_2_ monitor/controller and chamber, Biospherix). O_2_ and CO_2_ concentration was monitored continuously, such that their concentrations remain at 10% and 0.1% respectively. Exposure to normal air was limited to only during water, food and cage changes. All mice were sacrificed after 28 days (4 weeks) of exposure for analysis.

### Bone marrow chimeric mice generation

10-week-old recipient CD45.1^+^ and STING^-/-^ male mice were irradiated with two doses of 5Gy (11 minutes, 4-hour interval), and received bone marrow cells from 10-week-old CD45.2^+^ or STING^-/-^ donor male mice the next day through retro-orbital injection. Antibiotic (TMS, 100mg/kg) was delivered in drinking water, together with soft food for 2 weeks after bone marrow transplantation. Mice were given 6 weeks for bone marrow reconstitution before subjected to 4 weeks of chronic hypoxia exposure. An experimental design scheme is provided in **Figure 5A**.

### Sugen Hypoxia mouse model

10-week-old C75BL/6J and STING^-/-^ male mice were injected subcutaneously with SU5416 (Sigma Aldrich #S8442) suspended in a mixture of 0.5% carboxymethylcellulose sodium (Sigma Aldrich C4888), 0.9% sodium chloride (Sigma Aldrich #7647145), 0.4% polysorbate 80 (Sigma Aldrich #9005656), and 0.9% benzyl alcohol (Sigma Aldrich #100516) in deionized water. Mice received injection once a week at the dose of 20mg/kg body weight, while being exposed to hypoxia for 4 weeks. Control mice were injected with same volume of vehicle. After 4 weeks exposure of hypoxia, mice were returned to normoxia for 1 week before euthanasia.

### Treatment with anti-PD-L1 antibody

8-10-week-old LysM-Cre^+/-^STING^fl/fl^ and LysM-Cre^-/-^STING^fl/fl^ male mice were given *i.p.* injections of 500mg anti-PD-L1 antibody (BioXCell #BE0101) or isotype control (BioXCell #BE0090) once a week for 4 weeks, concurrently with injection of bleomycin (0.018U/g twice a week) or PBS per designated groups. Animal experiments and maintenance were approved by the Institutional Animal Care and Use Committee of University of Florida (IACUC; Protocol 08702). Animal studies are reported in compliance with the ARRIVE guidelines (Kilkenny, Browne, Cuthill, Emerson, & Altman, 2010) and with recommendations made by the Journal of Experimental Medicine.

### Primer sequences in mice

See **Table 1** for primer seαuences in mice. SAVI’s tails were sent to Transnetyx for confirmation of appropriate genotype.

**Table 1:**
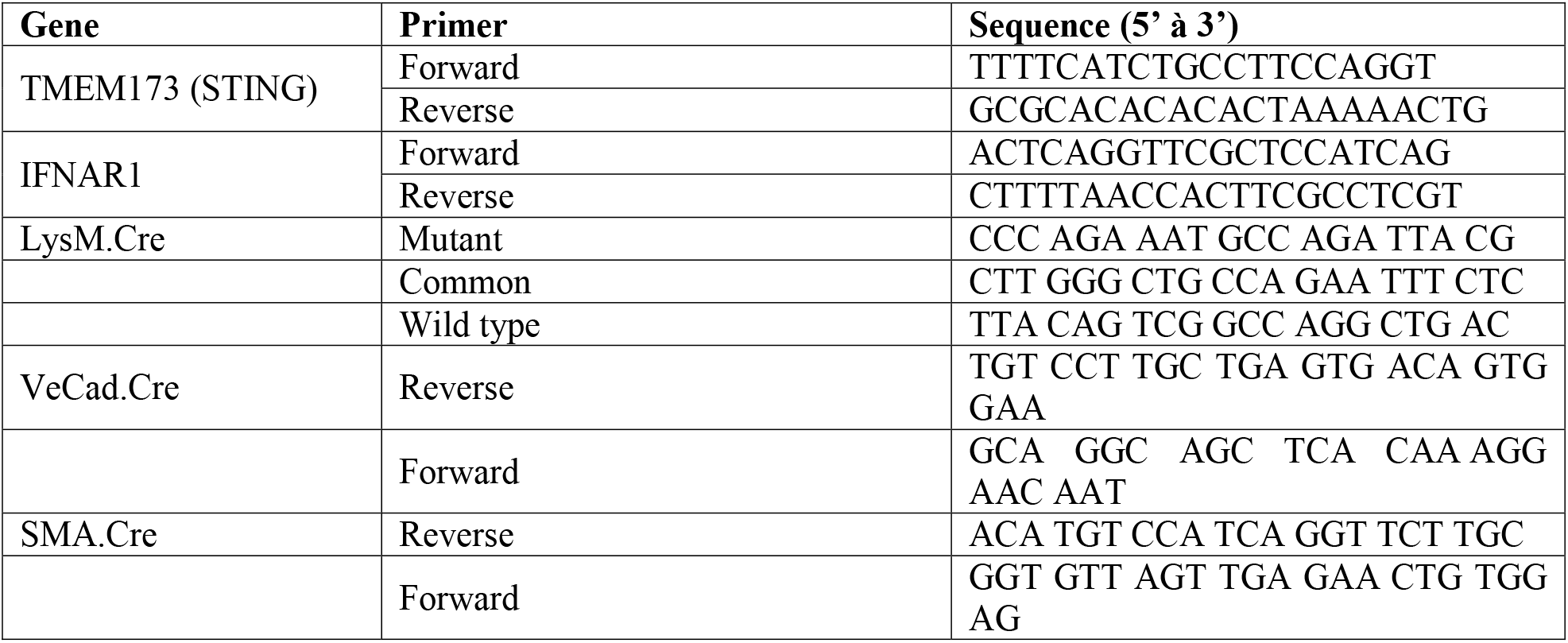
Primer sequences in mice

### Pulmonary hemodynamic assessment

Mice were put under deep anesthesia with intraperitoneal (*i.p.*) injection of 25% Avertin (2,2,2-tribromoethanol, Thermo Scientific^™^ #AC421432500) in PBS at 16mg/kg dose. A 1.4-French-pressure-volume microtip catheter (Millar Instruments, SPR-839) was inserted through a right internal jugular incision and threaded down into the right ventricle. The catheter was connected to a signal processor (PowerLab and ADInstruments) and the RVSP (mmHg) was recorded digitally and displayed with Chart5. After stable measurements of minimum 5 minutes, animals were euthanized with removal of hearts and lungs for subsequent analysis. The right ventricular was separated from the heart after removal of the atria, to which the weights of both right ventricle (RV) and left ventricle plus septum (LV+S) were obtained. Right ventricular hypertrophy was later calculated using RV/LV+S (%) ratio.

### Flow cytometry

Mouse’s left lung was cut into pieces and digested for 1h with 10mL of DNAse I (10mg/mL, Roche #10104159001) and type I collagenase (100mg/mL, Roche #11088882001) in PBS 10% FBS (Gibco #A4766801) at 37°C at 200rpm on a shaker for 1 hour. Digested cells were filtered through a 70mM cell strainer, and red blood cells were lysed using ammonium chloride lysis buffer (KD Medical #50-1019080). Single-cell suspensions obtained after using a 70mM cell strainer were stained with fluorochrome-conjugated surface antibodies for 30 minutes on ice, Fixable Viability Dye (eBioscience #65-0865-14) for 30 minutes or overnight on ice, fixed, permeabilized, and then incubated with intracellular markers for 30 minutes on ice. Data was acquired using BD FACSymphony A3 Cytometer (BD Biosciences) with 5 lasers and analyzed with FlowJo version 10 software.

### FACS antibodies

A comprehensive list of FACS antibodies used is presented in **Table 2**

**Table 2:**
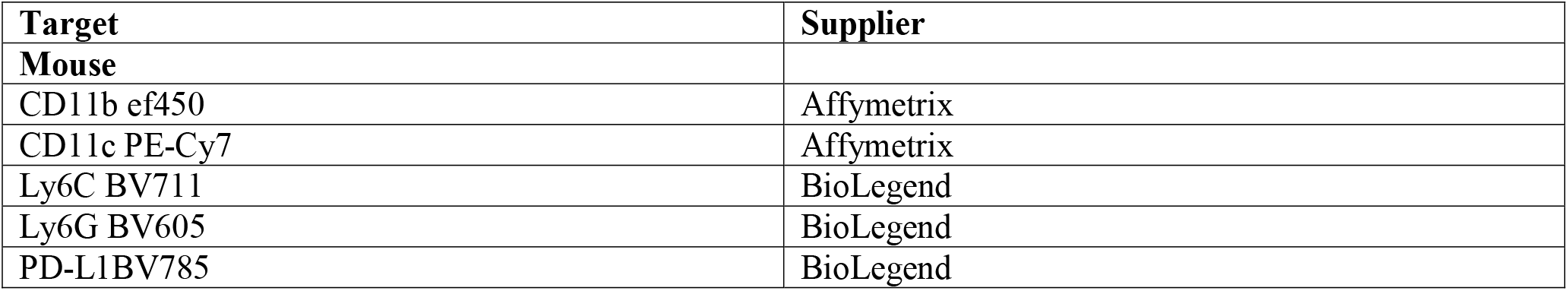
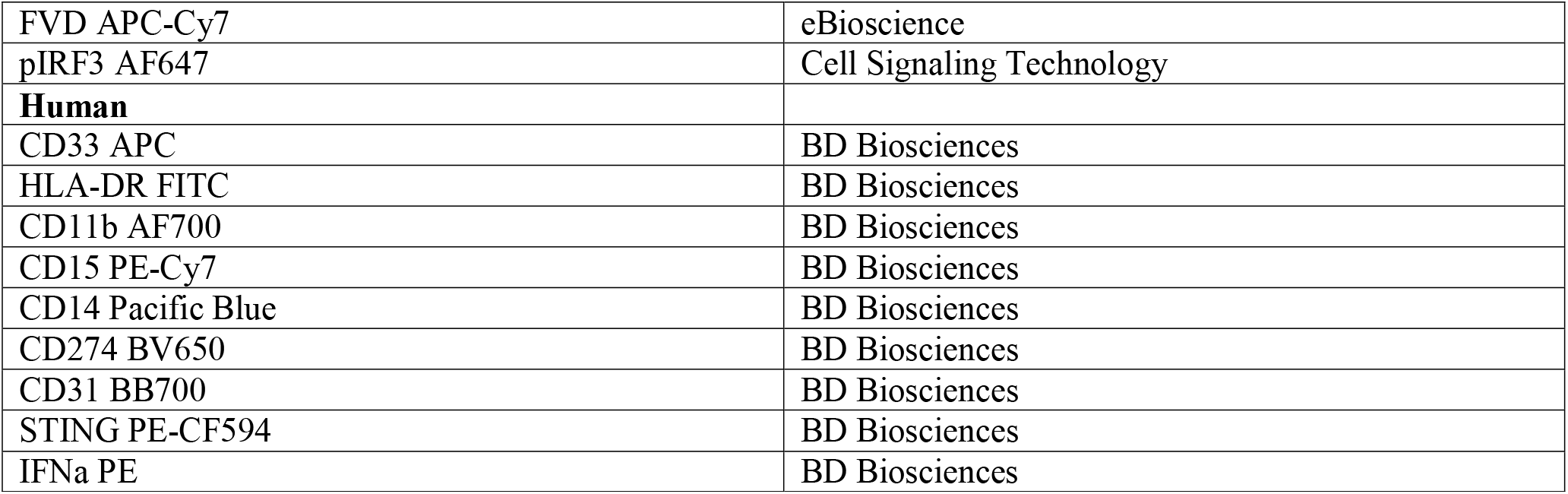
FACS antibodies for flow cytometry used in human and mice

| Target | Supplier |
| --- | --- |
| **Mouse** |  |
| CD11b ef450 | Affymetrix |
| CD11c PE-Cy7 | Affymetrix |
| Ly6C BV711 | BioLegend |
| Ly6G BV605 | BioLegend |
| PD-L1BV785 | BioLegend |
| FVD APC-Cy7 | eBioscience |
| pIRF3 AF647 | Cell Signaling Technology |
| **Human** |  |
| CD33 APC | BD Biosciences |
| HLA-DR FITC | BD Biosciences |
| CD11b AF700 | BD Biosciences |
| CD15 PE-Cy7 | BD Biosciences |
| CD14 Pacific Blue | BD Biosciences |
| CD274 BV650 | BD Biosciences |
| CD31 BB700 | BD Biosciences |
| STING PE-CF594 | BD Biosciences |
| IFNa PE | BD Biosciences |

### cDNA library construction and single-cell RNA-seq

Mouse whole lungs were perfused with 10mL PBS to remove blood cells. Lung tissues then were cut into small pieces and incubated in 3mL RPMI 1640 with 10% FBS (Gibco #A4766801), DNAse I (10mg/mL, Roche #10104159001) and 5% Liberase (Sigma Aldrich #05401127001) for 1h at 100rpm and 37°C. Tissues were triturated with 3mL syringes and 18G needles until complete dissociation. Cells are filtered through 70mm cell strainer, washed 3 times with DPBS 2% FBS and 1M EDTA to remove debris. Remaining red blood cells were lysed with ammonium chloride lysis buffer (KD Medical #50-1019080). Single cells were captured in the 10X Genomic Chromium Single Cell 3’ Solution, and RNAseq libraries were prepared following manufacture’s protocol (10X Genomics). The libraries were subjected to high-throughput sequencing on an Illumina NovaSeq 6000 platform, targeting 6000-8000 cells per sample with a sequencing depth of at 20 million reads of 150-bp paired-end reads.

### Process and quality control of the single-cell RNA-seq data

The raw sequencing reads were aligned with mouse genome mm10 provided on CellRanger website by 10X Genomics. The mapped reads then were used for unique molecular identifier (UMI) counting, following standard CellRanger pipeline for quality control as recommended by manufacture (10X Genomics). In short, cells with UMI counts lower than 500 or a feature count less than 200 were excluded. In addition, cells with greater than 30% of RNA content made up of the most common genes were excluded, as they accounted for empty droplets with free-floating RNA. Cells with greater than 30% of RNA content mapped to the mitochondrial genome were also eliminated as they indicated poor quality. Lastly, cells with abnormal high counts were discarded. Subsequently, the filtered single cells were imported into R package “Seurat” (version 4.0) for clustering of data and calculating differential gene expression, follow standard pipeline per manufacture instruction (10X Genomics). FGSEA package with gene sets from Molecular Signature Database, enrichPlot, and clusterProfile were used for generating differential expression of type I IFN genes. Markers for different population of cells (stromal, endothelial, and myeloid) were obtained from previously described work ^70–74^.

### Mouse histological staining

Mouse’s lung right lower lobe, upon harvest, was fixed in formalin overnight. Fixed tissues were paraffin embedded, cut with a Leica RM2235 Microtome, and stained for Masson Trichrome and a-smooth muscle actin to assess inflammation as well as to identify muscularized pulmonary vessels ^32^.

Masson Trichrome Staining (MTC) and semi-quantitative inflammation scoring: lung inflammation was evaluated on trichrome-stained lung sections using a 0 to 4 scale, with a score of 0, normal lung architecture; 1, increased thickness of some (£50%) of interalveolar septa; 2, thickening of >50% of interalveolar septa without formation of fibrotic foci; 3, thickening of the interalveolar septa with formation of isolated fibrotic foci; and 4, formation of multiple fibrotic foci with total or subtotal distortion of parenchymal architecture. Blind evaluation was performed on 10 randomized sequential, nonoverlapping field (magnification 10x) of lung parenchyma from each specimen. The mean score for the 10 fields represented the score for each individual specimen.

a-Smooth muscle actin (aSMA) staining and muscularized vessel count: formalin-fixed lung section was stained for rabbit polyclonal aSMA (Abcam #ab5694; diluted 1:750 in antibody diluent reagent solution [Life Technologies], not reused; blocking reagent Background Sniper [Biocare]). Stained lung specimens are then randomized and blindly assessed for pulmonary vessel counts on 10 sequential, nonoverlapping field (magnification 10x) lung parenchyma from each specimen. Muscularized pulmonary vessel is visualized in brown, either partially or completely. Vessel is considered small if £50mM, medium if 50-150mM, or large if >150mM.

### Image processing and acquisition

Images of representative tissues from histological staining were taken with Keyence BZ-X microscope at 10x magnification. Image processing was performed using BZ-X-Analyzer software (Keyence).

### T cell suppression assay

Wild-type C57BL/6 splenic-T cells are isolated using Stemcell EasySep^™^ Mouse T-cell Isolation kit (Stemcell #19851) and stained with Cell Trace Violet (ThermoFisher #C34571). These cells are then stimulated with anti-CD3/CD28 beads (Gibco #11456D) and coculture with splenic MDSC isolated using Stemcell EasySep^™^ Mouse Isolation kit (Stemcell #19867) at 3 different ratios (MDSC:T 1:1, 2:1 and 4:1). Cells are incubated for 4 days then T cells are sorted with flow cytometry using the following markers: CD4/CD8/CD25/Foxp3/Ror(g)t. Percent suppression is calculated as {[1- (proliferation with MDSC/proliferation without MDSC]x100} for each perspective group.

### Human samples

Paraffin embedded human lung tissues and single cell suspension of human lung digest were obtained from the Lung Research Data and Tissue Bank Registry at University of Florida. Ethical approval of biobanking and data collection was received from University of Florida Institutional Review Board (IRB201501114).

Single cell suspension of lung tissues was obtained from either healthy individuals (donor) or patients diagnosed with ILD through a lung transplantation (recipient). Samples were then processed by a specialist into single-cell suspension and stored at −80°C until staining for flow cytometry. Extracellular staining was performed with fluorochrome-conjugated surface antibodies for 30 minutes on ice, followed by Fixable Viability Dye (eBioscience #65-0865-14) for 30 minutes on ice. The cells were then fixed, permeabilized, and then incubated with intracellular markers for 30 minutes on ice. Data was acquired using BD FACSymphony A3 Cytometer (BD Biosciences) with 5 lasers and analyzed with FlowJo version 10 software.

Paraffin embedded lung tissues were cut with a Leica RM2235 Microtome and stained for rabbit polyclonal STING/TMEM173 antibody (Novus Biologicals #NBP2-24583SS; diluted 1:100 in PBS 10% BSA, not reused), followed by goat anti-rabbit IgG H&L (Abcam #ab6721). Protein of interest is visualized in brown, using DAB Substrate kit (Abcam #**ab64238**).

### Cytokine/Chemokine analysis

32 analytes (cytokines, chemokines, growth factors) in mouse whole lung protein were quantified using EMD Millipore’s MILLIPLEX Magnetic Bead Panel (Millipore Sigma #MCYTOMAG-70K), including: Eotaxin, G-CSF, GM-CSF, IFNg, IL-1a, IL-1b, IL-2, IL-3, IL-4, IL-5, IL-6, IL-7, IL-8, IL-9, IL-10, IL-12 (p40), IL-12 (p70), IL-13, IL-15, IL-17, IP-10, KC, LIF, LIX, MCP-1, M-CSF, MIG, MIP-1a, MIP-1b, MIP-2, RANTES, TNFa, and VEGF.

### Type I IFN ProQuantum assay

Serum collected from mice were analyzed for type I IFNa and IFNb using Mouse IFN ProQuantum Immunoassay Kit (Invitrogen #A46736 and #A47435), following manufacture’s protocols.

### Statistical analysis

Correlation analysis was performed using unpaired two-tailed Student’s t-test with GraphPad Prism 10. A p-value of <0.05 was considered statistically significant.

## Supporting information

Graphical abstract

## Figure legends

**Figure S1.**
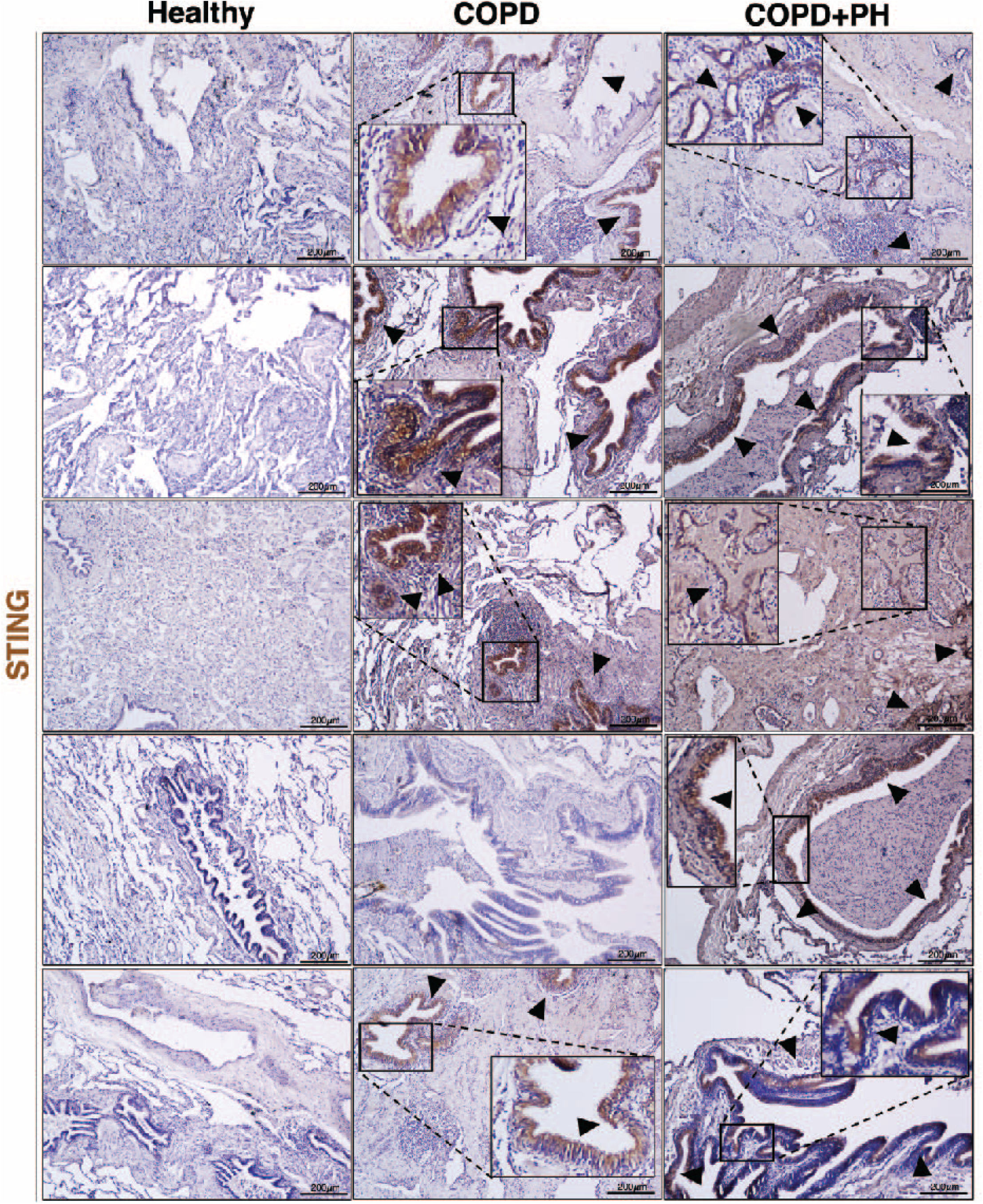
Patients with PH-predisposing lung disease display increased pulmonary STING expression. Representative images of IHC staining of STING (brown, arrowheads) in formalin-fixed lung sections of healthy individuals and patients with chronic obstructive pulmonary disorder (COPD) with or without PH at 10x magnification. Scale bar = 200mm. Each image is from an individual donor (n=5/group).

**Figure S2.**
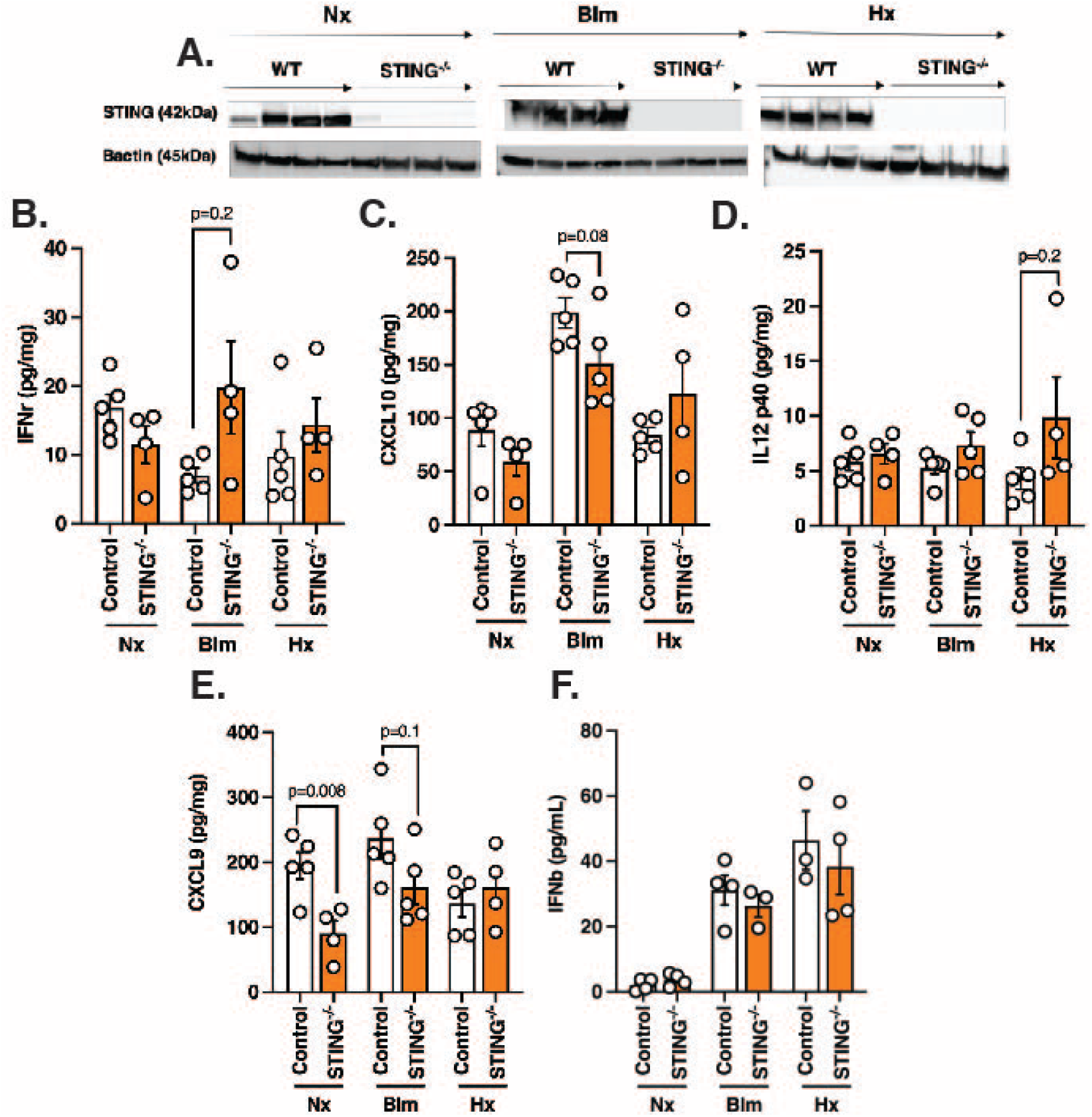
Global STING deficient mice are protected against PH development secondary to bleomycin and chronic hypoxia. (**A**) Western blot plot of STING expression in WT and STING^-/-^ mice exposed to different conditions. (**B – E**) Expression levels of (**B**) IFNr, (**C**) CXCL10, (**D**) IL12 p40 and (**E**) CXCL9 in lungs of designated groups. (**F**) Serum IFNb level of WT and STING^-/-^ mice. Each dot is an individual mouse (n=4-5/group). Column represents mean ± SEM. Significance level was calculated with un-paired two-tailed student T’s test. P<0.05 was considered significant.

**Figure S3.**
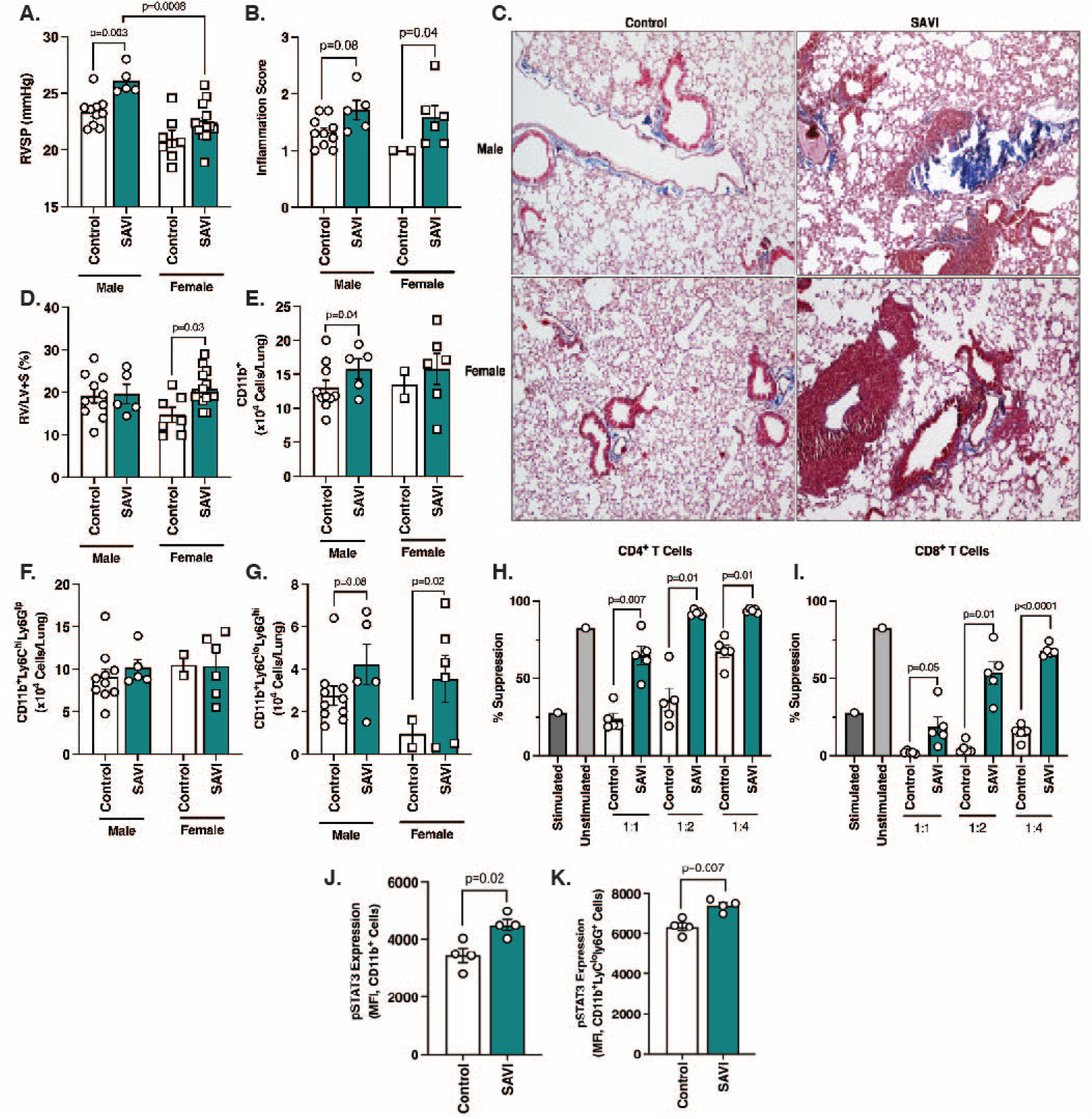
SAVI mice are genetically predisposed to pulmonary hypertension. (**A**) RVSP measurement of SAVI mice at baseline. (**B**) Quantification of inflammation degree by scoring MTC stained lung sections. (**C**) Representative images of MTC formalin fixed lung sections from the dedicated mouse groups at 10x magnification. Scale bare = 200mm (**D**) Fulton Index of SAVI mice at baseline. (**E** – **G**) Flow cytometric quantification of (**E**) CD11b^+^, (**F**) CD11b^+^Ly6C^hi^L6G^-^, and (**G**) CD1 1b^+^Ly6C^lo^Ly6G^+^ cells from lungs of designated groups. (**H and I**) Quantification of suppression capability of splenic MDSC isolated from control and SAVI male mice (n=5/group) on (**H**) CD4^+^ and (**I**) CD8^+^ T cells. % Suppression is measured by proliferation of T cells labeled with Cell Trace Violet (CTV) and stimulated with anti-CD3/CD28 antibodies at listed ratios. (**K and L**) Flow cytometric mean fluorescence intensity (MFI) quantification of pSTAT3 expression on (**H**) CD11b^+^Ly6C^hi^L6G^-^, and (**I**) CD11b^+^Ly6C^lo^Ly6G^+^ cells from lungs of SAVI male mice. Each dot is an individual mouse (n=4-10/group). Column represents mean ± SEM. Significance level was calculated with unpaired two-tailed student T’s test. P<0.05 was considered significant.

**Figure S4.**
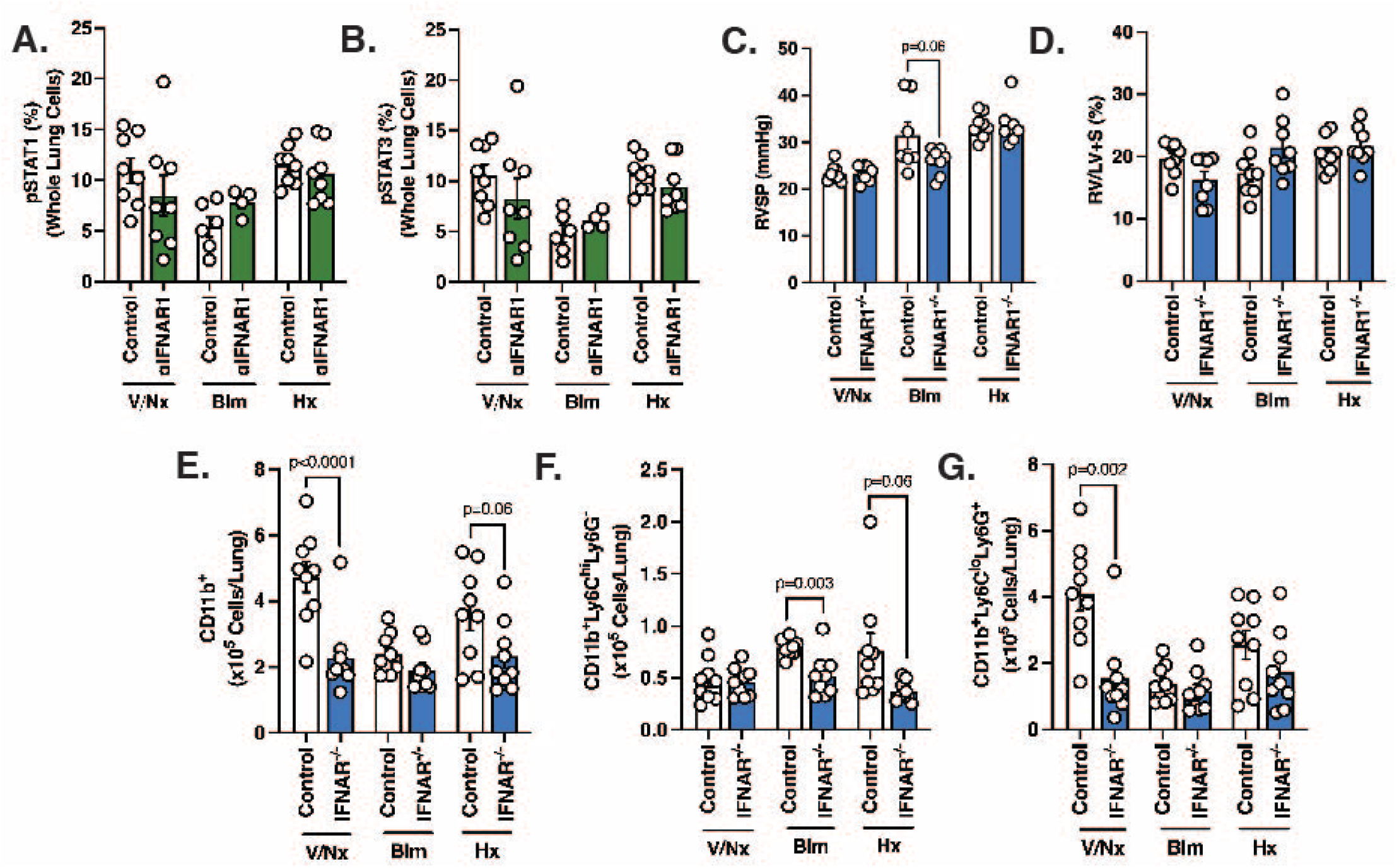
Type I IFN signaling plays minor role in PH development secondary to bleomycin and chronic hypoxia. (**A and B**) % Expression of (**A**) pSTAT1 and (**B**) pSTAT3 on whole lung cells of C57BL/6 mice from designated groups. (**C**) RVSP measurement of control and IFNAR1-/- mice subjected to normoxia, bleomycin, or hypoxia. (**D**) Fulton Index of mice from dedicated groups. (**E – G**) Flow cytometric quantification of (**E**) CD11b^+^, (**F**) CD11b^+^Ly6C^hi^L6G^+^, and (**G**) CD11b^+^Ly6loLy6G^+^ cells. Each dot is an individual mouse (n=3-8/group). Column represents mean ± SEM. Significance level was calculated with un-paired two-tailed student T’s test. P<0.05 was considered significant.

**Figure S5.**
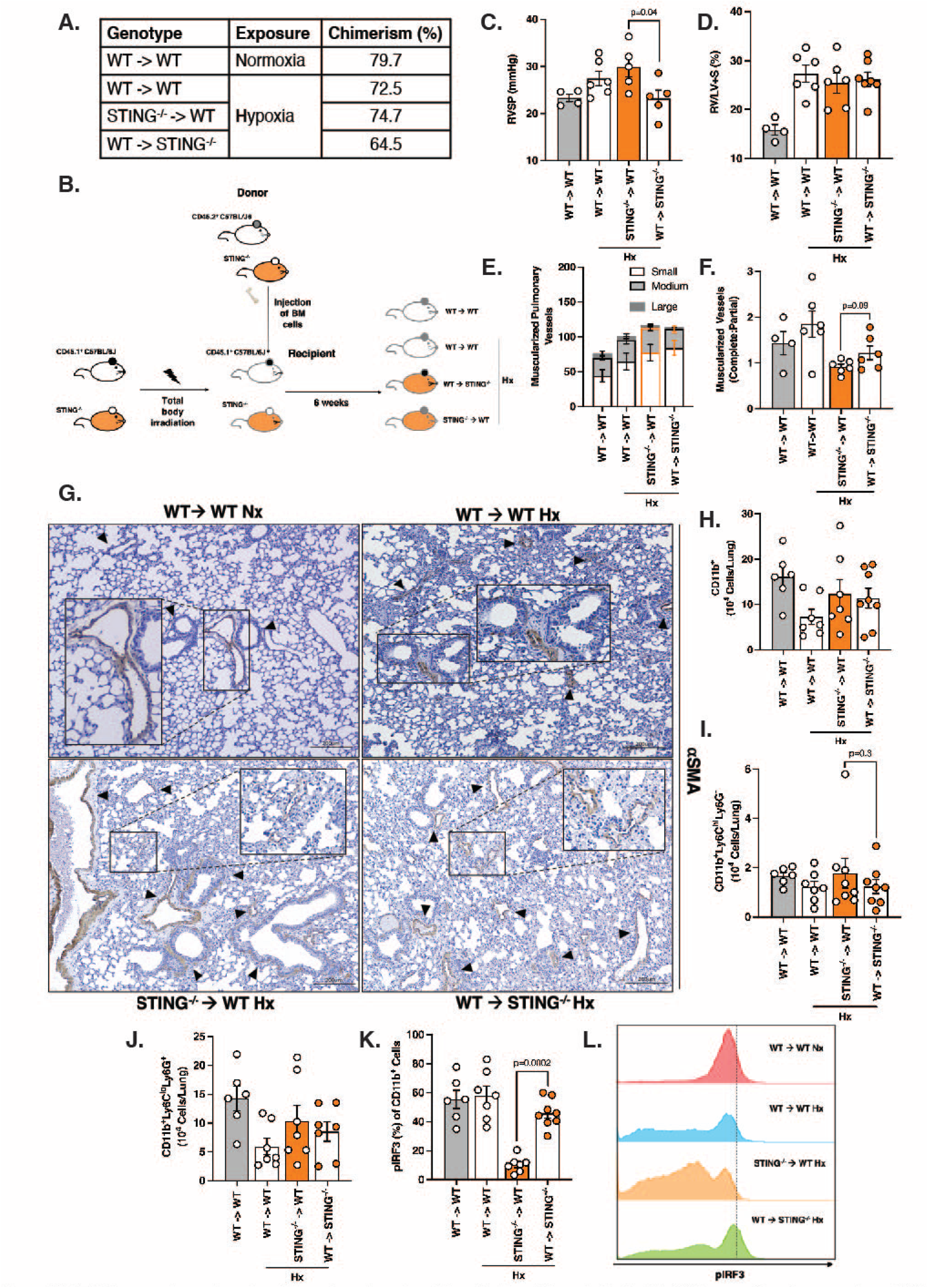
STING expression on hematopoietic and non-hematopoietic cells plays different roles in PH. (**A**) Average chimerism percentage of WT and STING^-/-^ chimeric mice in designated experimental groups. (**B**) Chimeric schematic to generate WT and STING^-/-^ chimeric mice. (**C**) RVSP measurement of WT and STING^-/-^ chimeric mice exposed to normoxia and chronic hypoxia (Hx). (**D**) Fulton Index of chimeric mice in dedicated experimental groups. (**E and F**) Quantification of small, medium, large, as well as complete and partial muscularized pulmonary vessels of experimental chimeric mice, assessing through aSMA IHC staining. (**G**) Representative images of aSMA (brown, arrowheads) IHC staining of formalin-fixed lung sections from chimeric mice of designated experimental groups. Scale bar = 200mm at 10x magnification. (**H** – **J**) Flow cytometric quantification of pulmonary infiltrated (**H**) CD11b^+^, (**I**) CD1 1b^+^Ly6C^hi^L6G^-^, and (**K**) CD1 1b^+^Ly6C^lo^Ly6G^+^ cells from designated mouse groups. (**L**) % Expression and (**M**) Flow representation of pIRF3 expression on CD11b^+^ cells from different experimental groups. Each dot represents an individual mouse (n=4-6/group). Column represents mean ± SEM. Significance level was calculated with un-paired two-tailed student T’s test. P<0.05 was considered significant.

**Figure S6.**
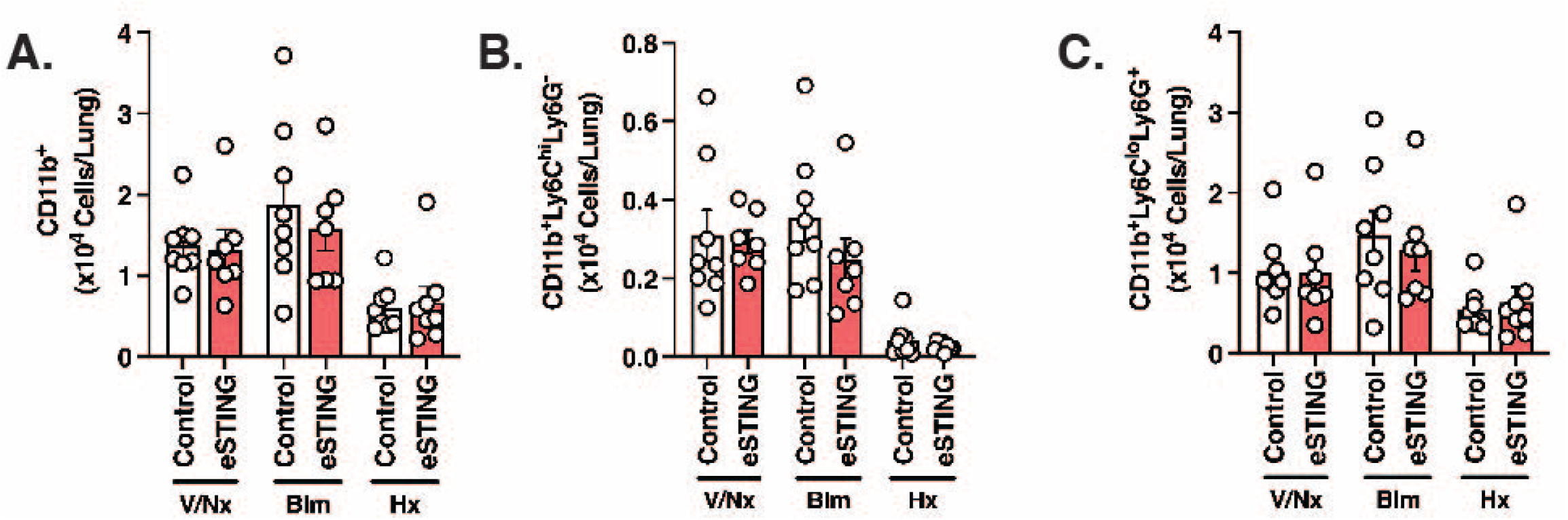
Smooth muscle, but not endothelial, specific deletion of STING provides protection against PH development secondary to chronic hypoxia exposure in mouse. (**A** – **C**) Flow cytometric quantification of pulmonary infiltrated (**A**) CD11b^+^, (**B**) CD11b^+^Ly6C^hi^L6G^-^, and (**C**) CD11b^+^Ly6^lo^Ly6G^+^ cells from designated mouse groups. Each dot is an individual mouse (n=8/group). Column represents mean ± SEM. Significance level was calculated with unpaired two-tailed student T’s test. P<0.05 was considered significant.

**Figure S7.**
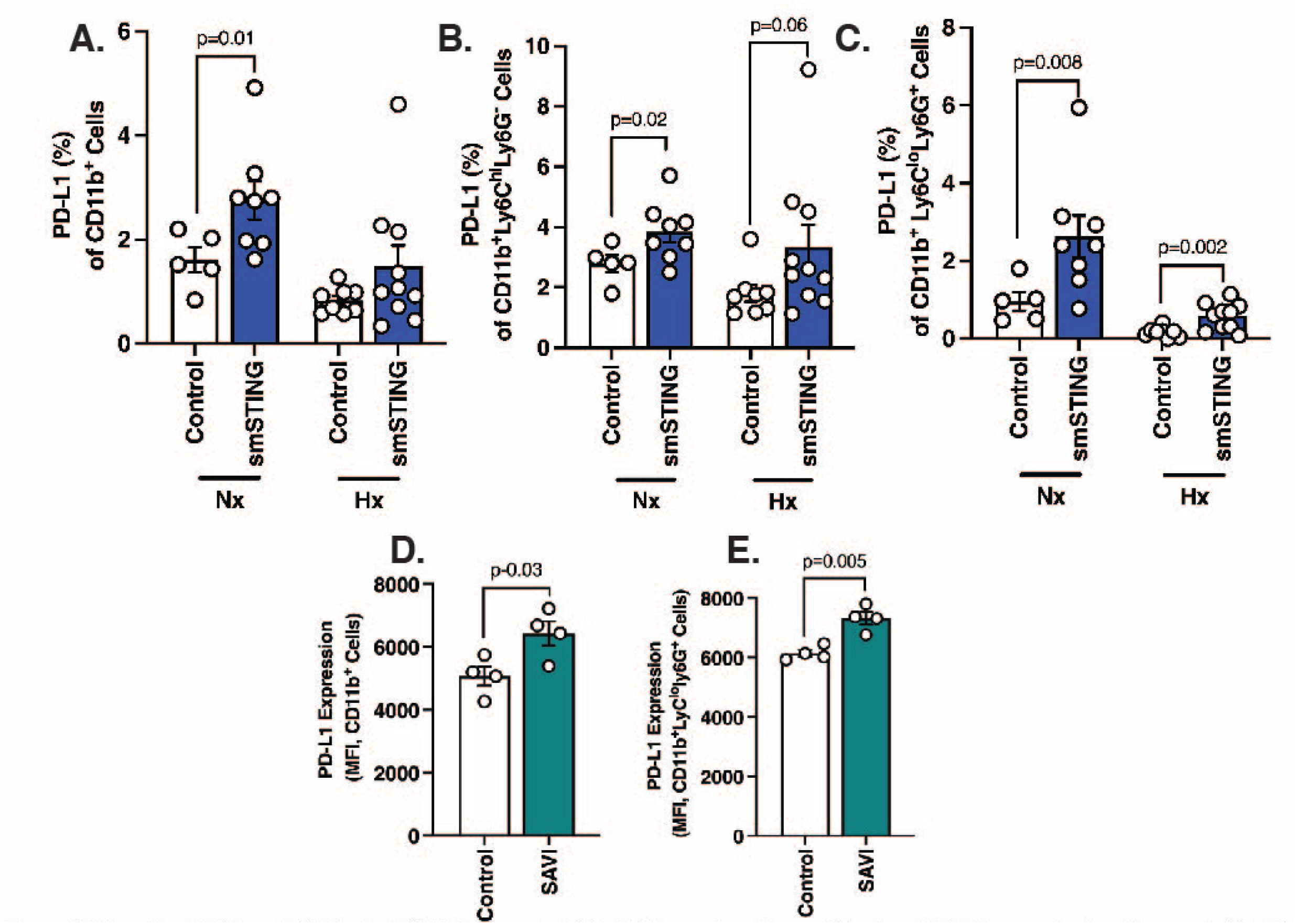
Severity of PH in mSTING mice is PD-L1 dependent. (**A** – **C**) Flow cytometric quantification of PD-L1 expression in pulmonary infiltrated (**A**) CD11b^+^, (**B**) CD11b^+^Ly6C^hi^L6G^-^, and (**C**) CD11b^+^Ly6C^lo^Ly6G^+^ cells of WT and smSTING mice in dedicated mouse groups. (**D and E**) MFI quantification of PD-L1 expression on (**D**) CD11b^+^ and (**E**) CD11b^+^Ly6C^lo^Ly6G^+^ cells of SAVI mice at baseline. Each dot represents an individual mouse (n=5-10/group). Column represents mean ± SEM. Significance level was calculated with un-paired two-tailed student T’s test. P<0.05 was considered significant.

## Supplemental figures

The supplemental material includes 7 figures describing (**S1**) IHC staining of STING in lung sections of healthy individuals and patients with chronic obstructive pulmonary disorder; (**S2**) western blot and of STING expression and expression level of different chemokines, cytokines and interferon in whole lung protein of WT and STING^-/-^ mice; (**S3**) PH assessment in SAVI mice; (**S4**) flow cytometric quantification of pSTAT1 and pSTAT3 in whole lung protein of mice treated with anti-IFNAR1 antibody, together with PH assessment of IFNAR1^-/-^ mice; (**S5**) PH assessment in WT and STING^-/-^ chimeric mice; (**S6**) flow cytometric quantification of infiltrated myeloid cells in eSTING mice; (**S7**) PD-L1 expression in infiltrated myleoid cells of smSTING mice and SAVI mice.

## Author contributions

A.T. Pham and A.J. Bryant wrote the manuscript, designed experiments, and performed data analysis. A.T. Pham, A.C. Oliveira, C. Fu, M.D. Alves, L. Mukhsinova, and E. Ebrahimi performed experiments. Z. Dupee analyzed scRNAseq data. A.T. Pham, M.D. Alves, H. Patel, R. Patel, and A. Nguyen performed histopathological analysis and scoring. L. Jin provided support and advised on experimental design. A.J. Bryant supervised the work.

## Acknowledgment

We thank Dr. Yuanquing Lu (University of Florida) for his generous help with RVSP measurement. We thank Dr. Matthew Schaller, Dr. Mark Brantly and Dr. Borna Mehrad (University of Florida) for generously sharing their equipment. We thank Dr. Jeffrey K. Harrison, faculties and trainees of University of Florida, Pulmonary Division for helpful discussion.

This project is supported by National Institute of Health (NIH) grants RO1HL142776, RO1HL142887 and University of Florida Gatorade Fund. A.C. Oliveira was supported by American Heart Association Postdoctoral Fellowship. The project was also supported by University of Florida Interdisciplinary Center for Biotechnology Research. The funders had no role in study design, data collection and analysis, decision to publish, or the preparation of the manuscript.

The authors declare no competing financial interests.

## Abbreviations

aSMA: Alpha smooth muscle actin
CTV: Cell trace violet
eSTING: VeCad-Cre^+/-^STING^fl/fl^ (endothelial specific deletion of STING)
IHC: Immunohistochemical
ILD: Interstitial lung disease
IPF: Idiopathic pulmonary fibrosis
IRF3: Interferon regulatory factor 3
MDSC: Myeloid derived suppressor cell
Mo-MDSC: Monocytic myeloid-derived suppressor cell
mSTING: LysM- Cre^+/-^STING^fl/fl^ (myeloid specific deletion of STING)
MTC: Masson trichrome
PAEC: Pulmonary arterial endothelial cell
PD-L1: Programmed death ligand-1
PH: Pulmonary hypertension
PMN-MDSC: Polymorphonuclear myeloid-derived suppressor cell
PVSMC: Pulmonary vascular smooth muscle cell
RVSP: Right ventricular systolic pressure
SAVI: STING-associated Vasculopathy onset in Infancy
SMC: Smooth muscle cell
smSTING: SMA-Cre^+/-^STING^fl/fl^ (smooth muscle specific deletion of STING)
STING: Stimulator of Interferon Genes
STING^-/-^: Global STING deficiency
VEGF: Vascular endothelial growth factor
WT: Wild type

