## Supplementary figures and images for "Cell dichotomous role of STING in pulmonary hypertension"

### Graphical abstract

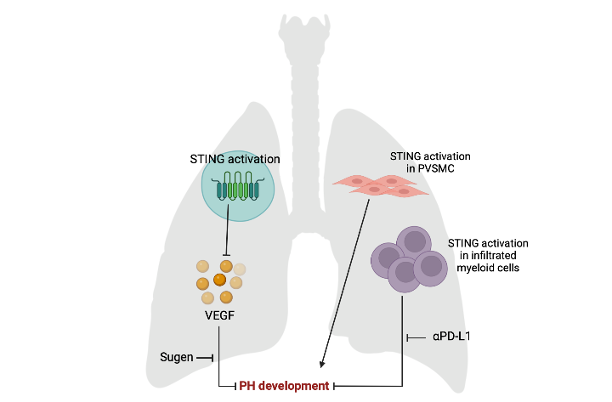
